# Transcranial direct and alternating current stimulation produce distinct long-lasting changes in macaque V1 stimulus-induced gamma

**DOI:** 10.64898/2026.05.18.726009

**Authors:** Niloy Maity, Supratim Ray

## Abstract

Transcranial direct or alternating current stimulation (tDCS or tACS) are used for the treatment of several cognitive disorders, many of which are due to imbalances in excitatory-inhibitory (E-I) interactions, but how stimulation affects the underlying cortical network remains an open question. In the primary visual cortex (V1), E-I interactions due to presentation of large gratings induce slow (20 Hz-35 Hz) and fast gamma (40 Hz-70 Hz) oscillations, which weaken with ageing and neurodegeneration and have been associated with different subtypes of interneurons. However, the effect of tDCS/tACS on stimulus-induced gamma is unknown. To investigate the impact of sustained stimulation on cortical E-I networks, we applied tDCS and tACS to two alert non-human primates while presenting full-screen gratings. We analyzed local field potentials before, during, and post-stimulation, focusing on gamma power and field-field coherency (FFC) as a measure of phase consistency. We found that tDCS significantly increased post-stimulation slow and fast gamma power, as well as FFC (60 Hz–100 Hz), with this effect lasting for approximately 1.5 hours. In contrast, tACS at 20 Hz consistently reduced slow gamma power, along with FFC (40 Hz–60 Hz). Our experimental observations were replicated in a physiologically realistic computational model of gamma generation by introducing targeted modifications to the synaptic weights within the simulated E-I network. Because long-lasting changes in local power and coherency strongly influence and modify cortical network, our findings reiterate the therapeutic and experimental potential of transcranial electrical stimulation to induce sustained modulation of cortical networks.

**SIGNIFICANCE STATEMENT:** Non-invasive transcranial current stimulation is used as a treatment for neural disorders, but how it modifies cortical dynamics to produce lasting changes is still debated. We recorded stimulus-induced gamma oscillations in macaque visual cortex as a readout of network interaction to study the effect of cortical stimulation. After stimulation over clinically relevant durations (∼20 minutes), direct and alternating current increased and suppressed gamma power and connectivity, respectively, and this effect persisted for more than an hour. We also found that modifying the synaptic weights informed by earlier research in a realistic gamma-generating model could replicate our experimental results. Our results establish gamma rhythm as a useful indicator to study the effect of neurostimulation on neural circuitry.

## INTRODUCTION

Stimulus-induced gamma oscillations in the primary visual cortex (V1) reflect network interactions between pyramidal neurons and inhibitory interneurons (1–3). In response to full screen gratings, this manifests as two distinct rhythms - fast (40 Hz-70 Hz) and slow gamma (20 Hz-35 Hz), which are tuned to distinct stimulus features such as size, contrast and spatial frequency (4–7) and are coherent over different spatial scales (8). In rodents, there is some evidence that they are mediated by parvalbumin (PV+) and somatostatin (SST+) subtypes of interneurons (1, 9, 10), although there are important differences between primate and rodent gamma (for example, rodents also have an extremely narrowband gamma around 60 Hz which is suppressed by visual stimulation and has feedforward thalamic origins (9–11). Importantly, perturbations in these oscillatory dynamics have been documented across a spectrum of neurodegenerative and neurodevelopment disorders, including Alzheimer’s Disease and Autism Spectrum Disorders (12, 13).

A promising strategy to restore or modulate such disrupted cortical dynamics is transcranial electrical stimulation (tES), a non-invasive neuromodulation technique that delivers low-intensity current through scalp electrodes to alter ongoing brain activity. Among tES modalities, transcranial direct current stimulation (tDCS) applies polarity-specific constant current, cathodal or anodal with the intent to bidirectionally regulate cortical excitability, and effects have been observed both during (online) and after (offline) stimulation (14–16). However, despite its mechanistic simplicity, the outcomes of tDCS have been far from linear (17) and have suffered from non-replicability (18), especially outside of the motor cortex.

Transcranial alternating current stimulation (tACS), by contrast, delivers oscillatory current at physiologically relevant frequencies, aiming to entrain, shift, or reinforce cortical rhythms. Accordingly, tACS has been shown to modulate cortical excitability (19, 20), entrain single unit spiking activity (21), enhance working memory (22), and show early therapeutic promise in cognitive disorders (23, 24).

While *in vitro* experiments, rodent studies, and computational models have explored the circuit-level effects of tES (25–27), how sustained current stimulation reshapes cortical network dynamics in the primate visual cortex, as reflected in visual-stimulus-induced gamma activity, remains largely uncharacterized. Both tDCS and tACS act primarily on the superficial cortical layers where pyramidal cell populations are densely concentrated, and stimulation of sufficient duration may engage synaptic modifications that outlast the stimulation period itself. Simulus induced gamma oscillations in V1, as a direct readout of pyramidal-interneuron circuit dynamics, may provide a principled probe to quantify stimulation-induced network changes.

Previous computational studies employed mean-field models to understand the oscillatory dynamics underlying excitatory-inhibitory (E-I) interactions in V1 circuits and successfully reproduced some features of gamma oscillations, such as their dependence on stimulus features (28, 29), stimulus discontinuities (30), shape (31) and temporal characteristics (32, 33). These models are governed by only a few parameters, allowing us to study how changes in synaptic weights within or between E and I populations affect gamma power.

We therefore first compared pre- and post-stimulation gamma power and field-field coherency to observe changes in oscillations and connectivity, respectively. Then, drawing on previous hypotheses of synaptic changes imposed by tES, we simulated the effect of stimulation in a gamma-generating E-I network model to test whether stimulation-driven changes in synaptic weights are sufficient to reproduce the experimental observations.

## RESULTS

We applied two types of transcranial electrical stimulation, tDCS and tACS, to two non-human primates while they performed a passive fixation task while full-screen gratings of varying spatial frequency, orientation and contrast were presented. These stimuli generated two distinct gamma oscillations in the visual cortex, termed slow and fast gamma. Both gamma rhythms become faster with increasing contrast, although while power typically increases with contrast for fast gamma, it peaks at mid-level contrasts for slow gamma (see Fig. 3 and Fig. 3-1 of (8) for details). In this dataset as well, fast gamma was more prominent for 100% contrast stimuli while slow gamma was strongest for 25% contrast (Fig. S1), and gamma peak frequencies varied with contrast (the slow and fast gamma ranges used for analysis for different contrast levels are shown in Fig. S1). The slow gamma range overlaps with the traditional beta range, but we term this as slow gamma since it is induced in V1 when visual stimuli are presented, unlike the beta rhythm which is prominent in sensorimotor areas and is modulated by motor movements (34, 35) . These grating stimuli were shown in ∼20-minute protocols six times: Pre (Before stimulation), Stim (actual or sham stimulation), Post (just after stimulation), and three more with 10-minute gaps (Fig. 1A). In the Stim protocol, we stimulated the right primary visual cortex of the monkeys with low-intensity current pulse (Fig. 1B) for 20 minutes while they viewed the same gamma-inducing visual gratings as other protocols. Sham stimulation experiments involved only ramping up the current to the stimulation intensity and immediately ramping it down over a duration of 10 s, at the beginning and end of the protocol, with no stimulation in between, and were used as a control condition in our study. Within each protocol, visual stimuli were presented in trials of 4.8 second duration which started with 1 second of gray screen after which 3 stimuli were shown for 800 ms with 700 ms inter-stimulus intervals (Fig. 1C). The monkey was given a juice reward for maintaining fixation within 1.5 degrees of the fixation dot for the entire trial.

**Fig. 1:**
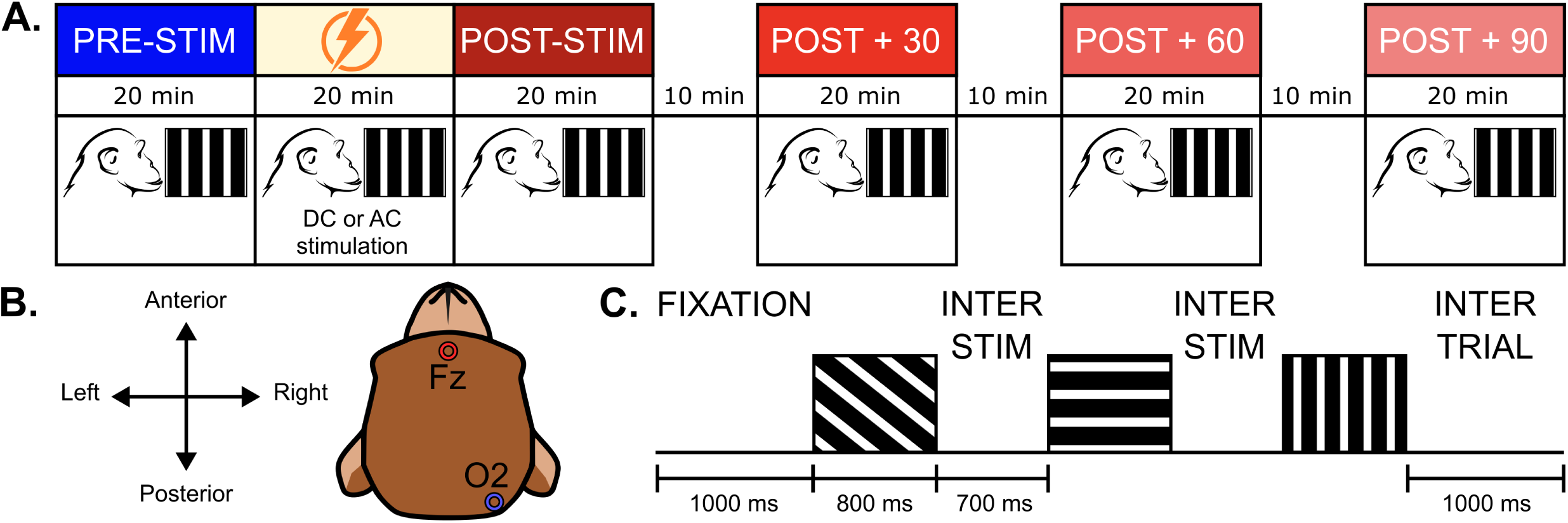
Experiment design and trial structure **(A)** Each experimental session consisted of six blocks of gamma protocol. Between the Pre-Stim and Post-stim block, stimulation was applied for 20 minutes. In case of Sham stimulation, current stimulation was replaced by only ramp up and ramp down of current for 10 seconds, in the beginning and end of gamma protocol. After post-stim block, three gamma protocols were recorded with 10 minutes of gap in between. **(B)** Stimulation electrode locations (Fz and O2) shown on a cartoon monkey head, the fronto-occipital montage targets right V1 where the Utah array was implanted. **(C)** Each trial started with a fixation dot, on successful fixation for 1 s, three stimuli displayed on screen consecutively for 800 ms each, with interstim gap of 700 ms. Example gratings of different spatial frequency, orientation and contrast are shown here. Each gamma block took ∼20 minutes to complete, with ∼140 trials.

### Cathodal tDCS enhances, whereas 20 Hz tACS suppresses Gamma power

The time-frequency representation of recorded data was visualized by averaging the spectrum across all trials and subtracting the baseline power from the stimulus-induced power (Fig. 2). In the tDCS experiments, data were collected from 160 LFP electrodes across five sessions (32 unique electrodes) for both the stimulation and sham experiments in M1. For M2, data were obtained from 118 and 119 LFP electrodes across three stim and sham experiment sessions, respectively (40 unique electrodes).

**Fig. 2.**
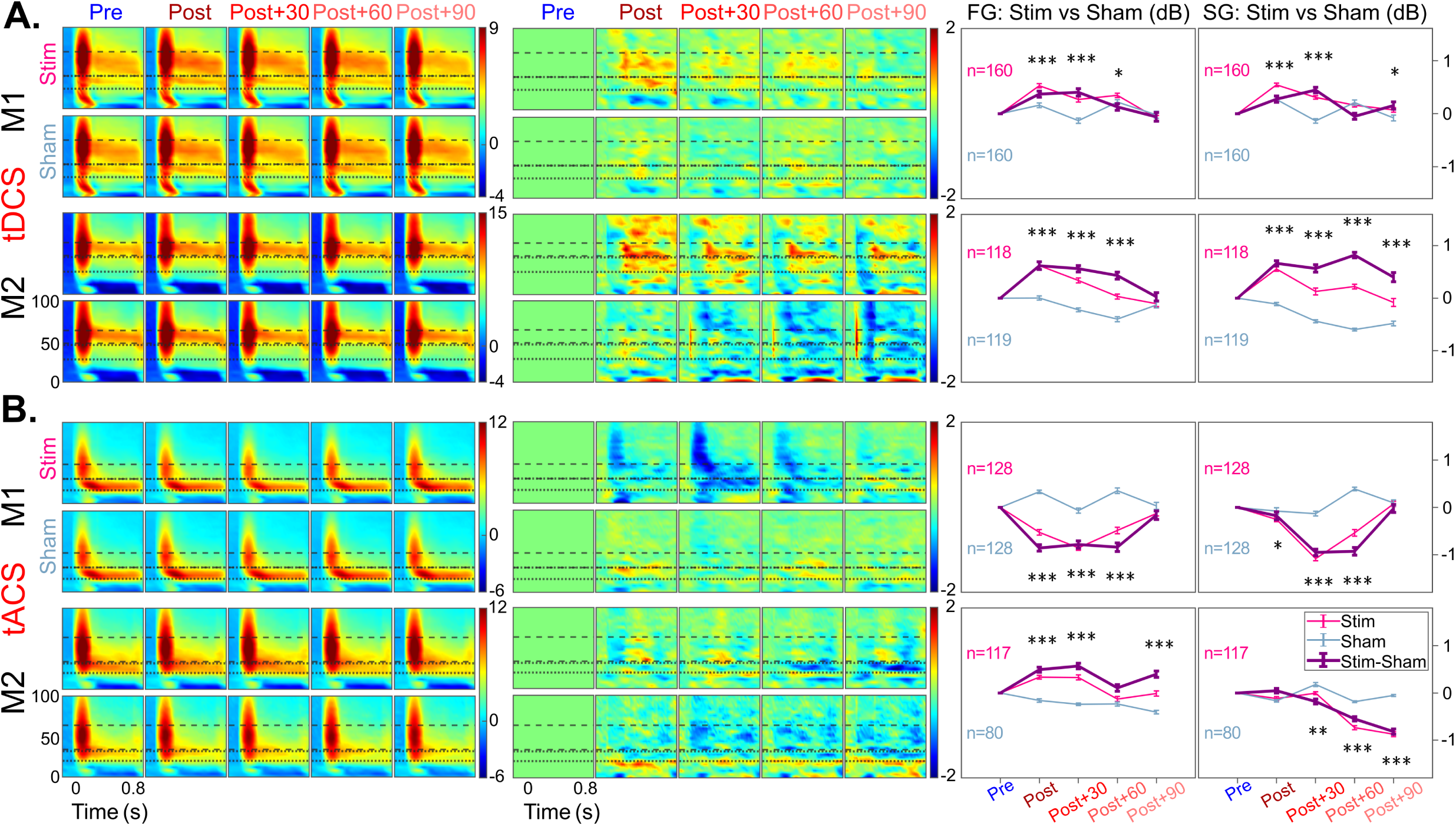
Effect of tDCS and tACS on stimulus-induced gamma power **(A , left columns)** Time frequency heatmaps for both stimulation and sham sessions are plotted for both monkeys in the cathodal tDCS condition, averaged across all electrodes pooled across all sessions. Stimulus onset is pointed as time ‘0’, and induced gamma was computed from 0.25 s to 0.75 s time range. Each protocol after post-stimulation had 10-minutes of gap between them. **(A, middle columns)** Pre-stimulation time frequency spectrums were subtracted from all the spectra from each protocol and plotted here, following the same structure of subplots in the left columns. **(A, right columns)** Line plots depicting change in gamma power, where pre-stimulation gamma band power is subtracted from all the protocols. The ΔGamma band power is plotted as a thin line in light red for the stimulation session and light blue for the sham session, with each data point corresponding to a specific protocol. Two sample t-tests were conducted on the corresponding values comprising all the electrodes from the stim and sham sessions, and significance is indicated with stars above the protocol (*p<0.05, **p<0.005, ***p<0.0005). Number of electrodes is mentioned in each subplot. The purple line is plotted after subtracting the sham session gamma band power from those of the stimulation session. **(B)** Same as **(A)**, but for 20 Hz tACS condition.

**Fig. 3.**
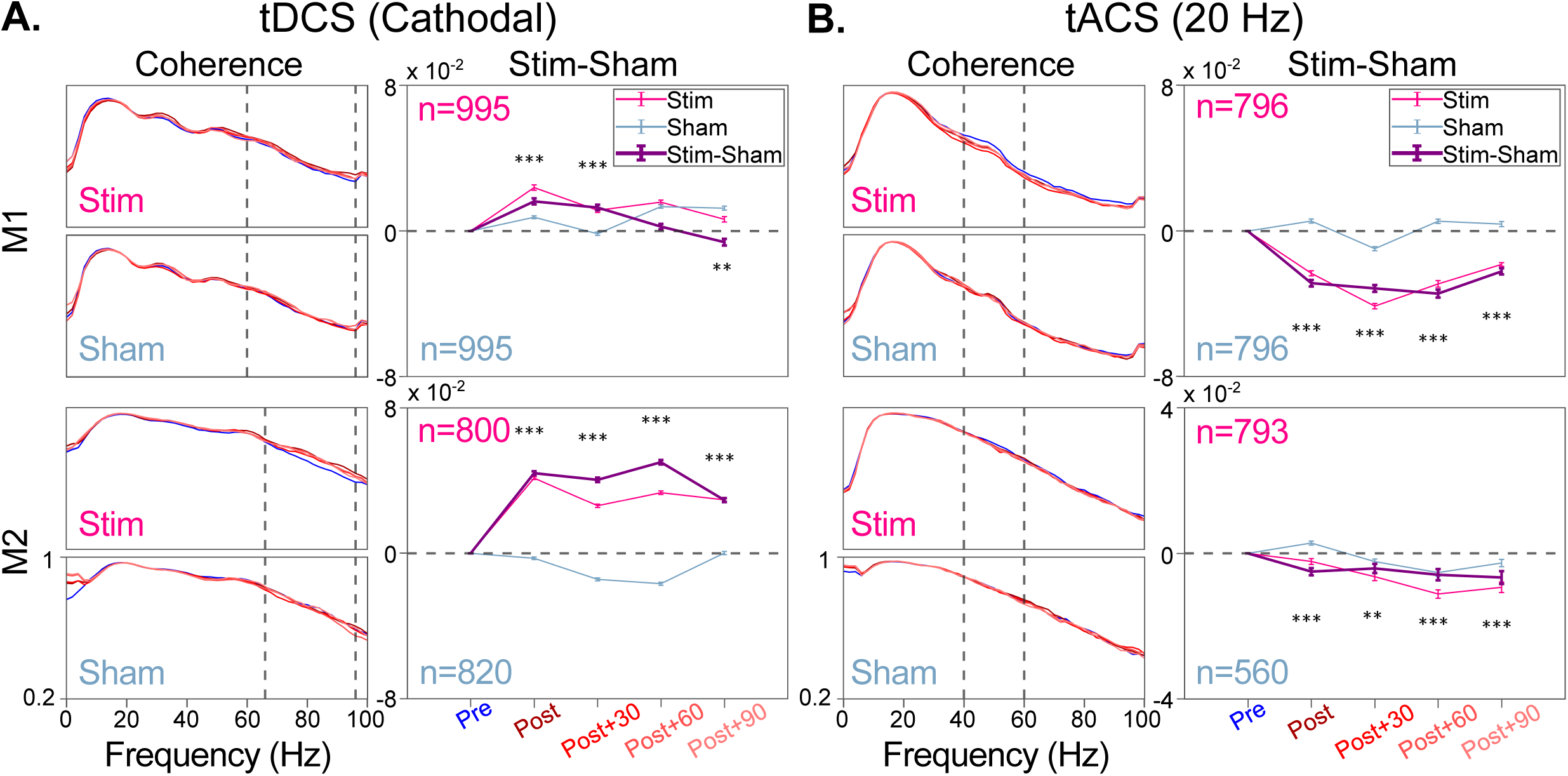
Effect of stimulation on LFP-LFP coherence **(A)** Average coherence values from all pairs with inter electrode distance in 0.4 mm -1.2 mm range. Different traces of red hue correspond to different post-stimulation protocols (left). Coherence values at selected frequency ranges (60 Hz-96 Hz for M1 and 66 Hz-96 Hz for M2) were averaged and plotted in line plots (right). **(B)** Same as **(A)**, but for 20 Hz tACS condition. Coherence values were averaged across 40 Hz-60 Hz range for both the monkeys and plotted in line plots (right).

Time-frequency spectra for the stimuli with 100% contrast showed that 3 mA cathodal tDCS increased power across the entire gamma range (Fig. 2A, left columns). This enhancement was evident in both the slow gamma band (24–40 Hz in M1; 28–46 Hz in M2) and the fast gamma band (40–70 Hz in M1; 48–64 Hz in M2) when compared to the pre-stimulus baseline. The effect was prominent across the following two post-stimulation protocols (Post, Post+30 min) in both monkeys. This change in oscillatory power tended to return to the pre-stimulation level by the last protocol (Post+90 min). To illustrate the change, we subtracted the entire change-in-power spectrum of each protocol with the change-in-power spectrum for the pre-stimulation condition (Fig. 2A, middle columns); since power is shown on a log scale throughout, this subtraction now shows the change-in-power relative to the power in the pre-stimulation protocol. This plot shows a clear increase in gamma power immediately after stimulation, which was observed for both gamma bands albeit appeared stronger in the fast gamma band, which persisted for about 1 hour. Gamma power relative to pre-stimulation condition for the stim condition (Fig. 2A, right columns; red traces) was significantly higher than sham condition (Fig. 2A, right columns; blue traces) for Post, Post+30 and Post+60 protocols (see Table S1 for details). We observed that in M2, overall power tended to decrease across blocks over time (also observed in other stimulation conditions for this monkey as shown below). However, the comparison relative to the sham was similar for both the monkeys (Fig. 2A, right columns; violet traces). For M2 also, the increase in gamma power for the Stim condition appeared to be stronger in the fast gamma range. However, as the suppression in the Sham condition occurred over both slow and fast gamma ranges, the difference between Stim and Sham conditions showed an increase in both gamma bands as shown in the right panels of Fig. 2A.

The change-in-power spectra shows the change in power relative to the pre-stimulus baseline, so we asked whether the pre-stimulus baseline power itself differed with stimulation. Fig. S2A shows the raw power spectral densities (PSDs) during baseline (-500 to 0 ms; dotted traces) and stimulus (250 – 750 ms; dashed traces) periods for different protocols (Pre, Post, etc.) and stimulation type (Stim and Sham). For M1, the baseline PSDs did not change across protocols (dotted lines were highly overlapping), while there was increase in raw power in both slow and fast gamma ranges post stimulation (dashed lines in blue were slightly below the reddish lines). For M2, there was some increase in power during the baseline period also (dotted blue line was slightly below other lines during both Stim and Sham conditions), in addition to the increase observed during the stimulus period (dashed blue trace was below the other traces). It is unclear why the baseline power increased over the course of the day for this monkey, which occurred in both Stim and Sham conditions and also for other stimulation paradigms such as tACS shown in S2B and discussed below). It could be due to changes in level of vigilance (36), or change in the noise level in the recording setup. Crucially, all comparisons were made between Stim and Sham (violet traces in the right panels of Fig. 2A), which at least partially accounted for the differences in baseline PSDs across protocols.

For 20 Hz tACS, the results were more variable across monkeys. In M1, we recorded from 128 LFP electrodes (across four sessions for both stimulation and sham experiments; 32 unique electrodes). In M2, data were collected from 117 LFP electrodes from three sessions of stim and 80 electrodes from two sessions of the sham experiment (40 unique electrodes). We applied ±1.5 mA tACS to M1 and ±2.5 mA tACS to M2 at 20 Hz. Fig. 2B shows the time-frequency plots for trials with stimuli of 25% contrast, which produced strong slow gamma. At this contrast, slow and fast gamma appeared at slightly lower frequencies (SG: 16-30 Hz in M1; 20-32 Hz in M2; FG: 30–48 Hz in M1; 34–64 Hz in M2; see Fig. S1). We observed that M1 showed a broadband suppression of fast and slow gamma by tACS, the suppression was only limited to slow gamma in M2 (fast gamma showed a small increase instead). The suppressive effect also built up with a delay, peaking at Post+30 protocol for M1 but continued to get stronger until the last block in M2. This suppression caused by tACS was significantly different from that in the sham sessions in both monkeys (p <0.0005, two-sample t-test) (Table S1).

We used gratings of three different contrasts (25%, 50%, and 100%), and, along with cathodal tDCS and 20 Hz tACS, we applied 10 Hz and 40 Hz tACS in separate sessions. Fig. S3 shows the effect of stimulation on gamma power for all contrasts and each stimulation condition, revealing an overall trend of increased slow and fast gamma power with tDCS and decreased slow gamma power with tACS across different contrast conditions. The 40 Hz stimulation showed stronger suppression of gamma activity for low-contrast stimuli in both monkeys. For 10 Hz tACS, the effect was weaker in general and inconsistent across monkeys.

### Firing rate and high gamma activity are less susceptible to post-stimulation changes

Previous studies have mentioned that the post-stimulation firing rate remains largely unchanged in the case of both transcranial direct and alternating current stimulation (37, 38). From both the stim and sham sessions in M1, we collected multi-unit activity (MUA) from 58 electrodes for tDCS and 43 electrodes for tACS experiments. From M2, we obtained multi-unit activity from 51 and 48 electrodes from stim and sham sessions of tDCS, while for tACS, we got data from 37 and 18 electrodes from stim and sham sessions, respectively. For each electrode, we normalized the firing rates by dividing by the maximum firing rate across protocols to ensure difference of firing rates between electrodes do not influence the calculation, and then computed the change in normalized firing rate from the pre-stimulation condition. We found that firing rates in M1 were slightly suppressed irrespective of the stimulation condition, but firing rates in M2 remained largely unaffected (Fig. S4).

There was a difference in the way the MUA was recorded in the two monkeys. Generally, the MUA is recorded by estimating a threshold based on the noise level (5 and 4 times the standard deviation in the signal for M1 and M2, respectively). Since tES could also change this noise level, we decided to fix this threshold during the pre-stimulation condition and use the same threshold for other protocols in M1. However, we found that as the sessions progressed, the quality of MUA (as measured using SNR) worsened. For M2, we therefore decided to recalculate the threshold before each protocol. To ensure that our firing rate results are not the consequence of difference in thresholding, we also calculated high gamma power (200 Hz-250 Hz) which is highly correlated with MUA (39) and does not depend on the choice of the threshold. We found that changes in high gamma power followed the same trend as firing rates for both types of stimulation (Fig. S4). Overall, compared to the changes in power in the slow and fast gamma ranges, the change in high gamma power was small and inconsistent across the two subjects (compare Fig. S4 with Fig. S3).

### Post-stimulation tDCS increases and 20 Hz tACS suppresses LFP-LFP coherence

To test the effect of tES on functional connectivity, we computed field-field coherence across electrode pairs separated by 0.4 (neighboring electrodes) to 1.2 mm (Fig. 3). Although the coherence traces were largely overlapping for different protocols, there was a small increase in coherence above 60 Hz (the blue trace corresponding to the pre-stimulation condition was below the red colour traces in both monkeys in this frequency range). We averaged the coherency values (60 Hz-96 Hz in M1 and 66 Hz-96 Hz in M2) for each protocol and subtracted the pre-stimulation values, which revealed a significant increase in coherency post stimulation in both monkeys (Fig. 3A), consistent with the trends observed in power albeit the chosen frequency range was higher in this case.

Stimulation at 20 Hz tACS, on the contrary, lowered coherency compared to sham sessions over a broad frequency range above 40 Hz in M1 and within 40 Hz-60 Hz in M2 (Fig. 3B). The effect of tACS at 10 Hz and 40 Hz was weaker and more inconsistent across two monkeys (Fig. S5). Interestingly, these trends did not weaken even when electrode pairs with larger interelectrode distance were used (Fig. S6).

### Change in synaptic weights in model explains changes in gamma power

To investigate the underlying mechanism of the post-stimulation changes, we employed a computational E-I mean field model comprising excitatory and inhibitory populations (40). Following Jadi and Sejnowski’s work (29), we chose input value pairs for the excitatory and inhibitory drives (i_E_ and i_I_, respectively) in a superlinear regime, as a prerequisite for generating realistic gamma oscillations, and i_I_ > i_E_ to emulate our full-screen grating stimuli condition, which causes a strong surround-suppressing inhibitory drive (41). This was further tuned to represent fast and slow gamma oscillations (Fig. S7A; detailed description in materials and methods), as done recently (42). Here, fast and slow gamma were generated by two independent E-I populations, where fast gamma was due to Parvalbumin positive (PV+) interneurons while slow gamma was due to the Somatostatin (SOM) interneurons, although we note that this is based on rodent studies (9, 10), and gamma oscillations in rodent and primate gamma could be different (11). From the simulated oscillations, we extracted peak power from the relevant frequency range and tested how changing the synaptic weights affected gamma power.

We found that modulating the excitatory-excitatory synaptic weight (W_EE_) exerted the most profound influence on gamma power; specifically, changes in W_EE_ were directly and positively correlated with power shifts in both fast and slow gamma bands, with larger changes in fast gamma compared to slow (Fig. S7B). This is consistent with the analytic results obtained by Jadi and Sejnowski (see equation 6 and 7 in the Methods section). The peak frequencies also shift leftward (Fig. S8) or rightward (Fig S9) with increase or decrease in W_EE_, as predicted analytically. Although these analytic results do not predict any dependence on inhibitory-to-excitatory weights (W_EI_) or excitatory-to-inhibitory weights (W_IE_), we found a weak positive dependence of W_IE_ on fast gamma (i.e., increase in power with increase in weights) and a weak negative dependence of W_EI_ for slow gamma (Fig. S7B). Note that following the known physiology of the somatostatin network, there was no self-coupling of the inhibitory population (W_II_ set to zero for slow gamma). We also simulated changes in W_II_ for fast gamma and found a small negative dependence, as predicted analytically (data not shown). When these weights were adjusted in pairs (W_EE_ with W_EI_, and W_EE_ with W_IE_), the oscillatory output of the network remained primarily dictated by fluctuations in W_EE_ (Fig. S10A and S10B), identifying recurrent excitation as a critical factor for gamma modulation.

Our experimental results do not put sufficient constraints on these models, since we mainly observed a change in power (without any noticeable change in peak frequency), which could be explained by manipulating many different combination of weights. Nevertheless, if there are any specific mechanisms proposed in the literature, these models can be used to test those predictions. Thus, building on evidence that tDCS promotes dendritic spine growth and structural remodeling in pyramidal neurons (43–45), we simulated the “offline” effects of tDCS by incrementally increasing all three synaptic weights (W_EE_, W_IE_, and W_EI_) by 2%-6%. This global enhancement of synaptic efficacy resulted in a significant increase in fast gamma power (Fig. 4B, top left) and a modest increase in slow gamma power (Fig 4B, down left), mirroring our post-tDCS observations.

**Fig. 4.**
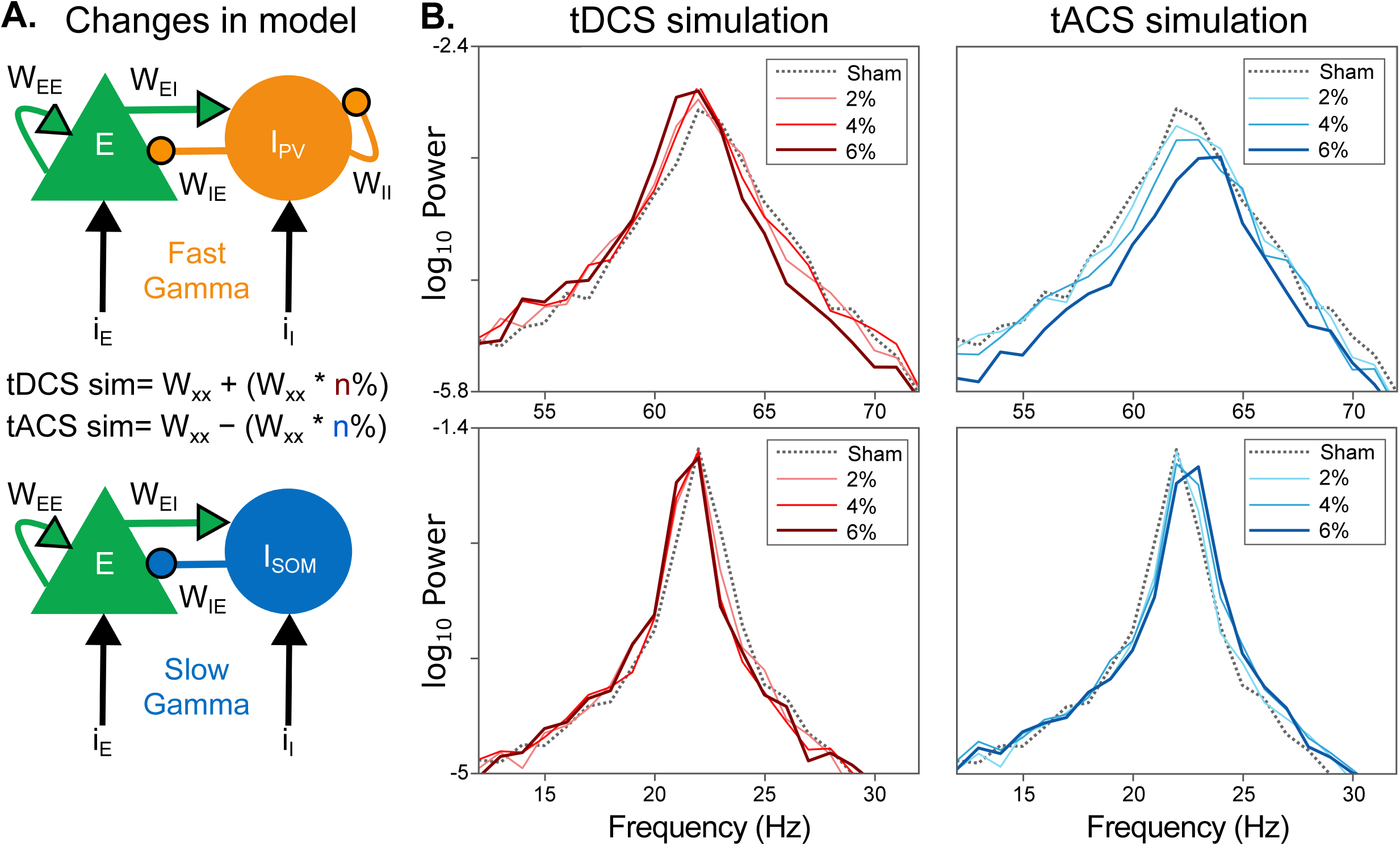
Change in synaptic weights simulate effect of current stimulation on pyramidal interneuron network gamma model **(A)** Change in model with increase in W_EE_, W_EI_ and W_IE_ as our hypothesis for tDCS condition based on (44, 45), and decrease in W_EE_, W_EI_ and W_IE_, for tACS condition as our hypothesis based on (46, 74). Model parameters for fast gamma and slow gamma conditions were used in simulation of PSD for both tDCS and tACS. **(B)** Change in PSD simulated in tDCS (left column) and tACS (right column) condition for both fast gamma and slow gamma with incremental change in synaptic weights.

To simulate the suppressive effects of tACS, we focused on the “frequency mismatch” hypothesis, which posits that a fixed-frequency external drive decouples neurons from the fluctuating endogenous peak frequency of the network (46). We modelled this loss of synchrony by implementing an incremental decrease (2%-6%) in the recurrent excitatory synaptic weights (W_EE_), reflecting a functional “detuning” of the pyramidal population, and a concurrent reduction in W_IE_ and W_EI_ as well as a detuned pyramidal network might disrupt the rhythmic input it sends to interneurons. This reconfiguration of synaptic weights resulted in a robust, dose-dependent decrease in gamma power mirroring our experimental observations (Fig. 4B, Right panels). Note that while both tDCS and tACS results can also be explained by only increasing or decreasing the excitatory weights (W_EE_) alone, this particular manipulation also changes the peak frequency substantially (Fig S8 and Fig S9). However, the peak frequency has opposite contributions from W_EE_ versus W_EI_ and W_IE_. Therefore, increasing all three concurrently effectively reduces the shifts in peak frequency, which better explains our results. Overall, these computational results suggest that the divergent effects of tDCS and tACS on cortical oscillations can be parsimoniously explained by opposing shifts in the efficacy of excitatory and inhibitory connections in the cortical network.

## DISCUSSION

We demonstrate for the first time in non-human primates that non-invasive neuro-stimulation induces lasting changes in V1 circuit dynamics that is reflected in stimulus-induced narrowband gamma power and field-field coherency. Cathodal tDCS enhances both gamma power and field-field Coherency; in contrast, gamma power is suppressed and coherency is reduced by 20 Hz tACS across recording sites. We hypothesized that these modulations arise from changes in synaptic weights within a gamma generating Excitatory-Inhibitory (E-I) network. By adjusting E-I weights in a network model based on prior stimulation studies, we found that the model output matches our experimental findings.

### Stimulus induced gamma as substrate of network state

Coordinated neural responses to a stimulus manifesting as oscillations are often task-dependent, brain state-dependent, or reflective of the macroscalar properties of a network. In the visual cortex, while fast gamma oscillations are primarily governed by the excitatory pyramidal cell (PYR) and inhibitory parvalbumin-positive (PV+) interneuron network, somatostatin-positive (SST+) interneurons mediate long-range horizontal connections with pyramidal cells, driving slow gamma activity (9, 10). Consistent with these frameworks, dysfunctional hyper- or hypo-plasticity within cortical gamma networks is linked to various cognitive deficits, including those observed in Schizophrenia and autism related disorders (47). Furthermore, grating-induced gamma oscillations are notably weaker in patients with Alzheimer’s disease than in healthy individuals (12).

### Caveats of non-invasive neurostimulation: methodological variability

Non-invasive brain stimulation (NIBS) techniques, such as tDCS and tACS, modulate both cortical surfaces (14, 19, 27, 48) and deeper structures (21, 49) depending on the targeted region and behavioural context. While cellular-level studies (21, 25, 26, 50) and diverse brain areas of various animal models (27, 37, 51, 52) have provided fundamental insights, understanding how cortical networks respond to sustained stimulation, and the resulting post-stimulation effects, remains a prerequisite for clinical translation.

A significant challenge in NIBS is the attenuation of current by the scalp and skull; a substantial portion of the applied electrical energy is dissipated before reaching the brain (53, 54). This raises questions regarding the optimal parameters required to drive meaningful, long-lasting neural changes. Non-human primates (NHPs) are uniquely suited to address these concerns; like humans, they possess thick skulls and gyrencephalic brains, providing an ideal model to validate non-invasive stimulation methods through invasive recordings of local field potentials (LFP) and multi-unit activity.

‘Offline’ or post-stimulation effects have been investigated across a wide range of intensities and durations in tDCS and tACS research. While the relationship between applied intensity and physiological effect is often characterized as dose-dependent (38, 55), it can be characteristically variable (17) and longer duration stimulations; typically ∼20 minutes are necessary to reset the brain activity into state of lasting post-stimulation effect (56–58). In our study, we opted for 3 mA cathodal tDCS, and 1.5 mA-2.5 mA tACS (at 20 Hz) for 20 minutes using HD-tES ring electrodes (59), to ensure that the current was delivered focally and at a tolerable dosage to elicit robust post-stimulation effects.

### tDCS and tACS modify synaptic weights in cortical network

Although traditional polarity-specific rules often characterize cathodal tDCS as inhibitory, higher intensities have been shown to increase cortical excitability (17) and these differences might be explainable by morphology of cortical neurons, as distal dendrites are expected to be strongly depolarized in cathodal stimulation (25, 60) as seen in, in vivo and modelling studies (61, 62), Such dendritic depolarization by stimulation can trigger calcium-mediated spine growth and structural remodeling (43–45). In cortical network, pyramidal-parvalbumin (PYR-PV+) interactions are heavily influenced by dendritic activity (63) and often they can operate without interference of soma (64). Furthermore, strong calcium mediated AMPAR activity and strengthening of PYR-PV+ connections has shown increase in gamma power in hippocampus (65).

An increase in dendritic spine density on pyramidal cells effectively expands the pool of available synaptic contact points for both recurrent pyramidal and interneuron populations, which could effectively increase the effective synaptic weights within the gamma-generating circuit. In our E-I model, a global increase in the weights of PYR-PYR (W_EE_), PYR-PV+ or PYR-SST+ (W_IE_), and PV+-PYR or SST+-PYR (W_EI_) connections led to a robust increase in gamma power, providing a plausible mechanism for the sustained post-stimulation enhancements observed in our data.

In a seminal non-human primate study, Krause et al. (37) have shown that during tDCS, field-field coherence (FFC) increases between brain regions in gamma ranges. We found that the changes in FFC occurred at a higher frequency (60-100 Hz) range compared to the frequencies where gamma power was observed or modulated by electrical stimulation. Power in the high-gamma range above ∼80 Hz has been shown to be tightly coupled with the aggregate firing of the population of neurons around the microelectrodes (39). In a cross-frequency-coupling study, slow gamma phase has been shown to be preferentially coupled to power between 80-150 Hz, while fast gamma was coupled to frequencies above 150 Hz (66). The slow gamma and 80-150 Hz coupling was particularly strong for electrocorticogram (ECoG) signals. This could potentially be explained by assuming that power between 80-150 Hz reflects dendritic spikes (which are typically broader than somatic spikes), as somatostatin interneurons are thought to target the dendrites, and dendritic activity is likely to be better picked up by superficial ECoG electrodes. Under this assumption, enhanced coupling of dendritic spikes could then reflect in enhanced FFC in our data. Another possibility is that this enhanced FFC simply reflects more synchronized firing, which is better captured in the lower frequency range (60-100 Hz) which are less susceptible to small jitters in spike timing compared to higher frequencies (67).

Regarding tACS, changes in oscillatory power at targeted frequencies have been widely reported in EEG studies, particularly within alpha band in visual cortex (58, 68), beta or gamma bands in motor cortex (69, 70). These ‘offline’ changes are thought to be driven by NMDA-dependent plasticity (71), and recently among major depressive disorder patients, alpha tACS resulted in decreased alpha power and improvement in symptoms (24). Though there are examples of tACS effectively causing changes lasting up to 70 minutes (58), how a network responds to these changes is unclear with claims of absence of post-stimulation effects in humans (72, 73) and non-human primates (38).

Recent studies in non-human primates have highlighted a paradoxical feature of tACS: while it can increase phase-locking value (PLV) of responding neurons without altering mean firing rates (21), stimulating already-entrained population at its dominant frequency can significantly decrease their PLV (46). This suggests that a strongly oscillating network may become “detuned” under rhythmic stimulation, with subsets of pyramidal neurons losing synchrony and driving a reduction in recurrent excitatory (W_EE_) synaptic weights. The authors also modelled this phenomenon and reasoned that it is likely rooted in the inherent variability of neural rhythms; because peak frequencies of oscillations slightly fluctuate across trials, a fixed frequency tACS protocol inevitably creates a frequency mismatch that degrades network synchrony ((46), see their Fig. 4). Supporting this mechanistic view, recent simulations of strongly coupled E-I networks demonstrate that tACS applied at the dominant frequency (35 Hz) results in a post-stimulation reduction of PYR-PYR synaptic weights and a concomitant drop in the power spectrum ((74), see their supplemental Fig. S7, middle row). Moreover, while some debate remains (75), evidence suggests that pyramidal neurons may be more susceptible to external electric fields than interneurons (25, 76), potentially making the recurrent excitatory loop particularly vulnerable to tACS-induced disruption. This proposed decrease in W_EE_ likely triggers a downstream decoupling of the PYR-SST+ or PYR-PV + and SST+-PYR or PV+-PYR feedback loops, causing reductions in W_IE_ and W_IE_ respectively in the gamma model.

In our study, we observed that 20 minutes of tACS suppressed gamma power and disrupted post-stimulation field-field coherence (FFC) in the 40–60 Hz range across both the monkeys. To investigate the underlying mechanism, we found effect of changing PYR-PYR synaptic weights (W_EE_) had the most profound effect on gamma power when all the weights (W_EE_, W_IE_ and W_EI_) subjected to same percentage of decrease. Taken together, these results suggest that tACS-induced gamma suppression is possibly mediated by a specific de-tuning of recurrent pyramidal networks.

Another reconciling finding is that firing rates remain unchanged after stimulation in both tDCS and tACS experiments. While applying tACS and tDCS, previous NHP studies reported no change in firing rates, even in comparable stimulation intensity (21, 38). This supports the notion that the observed network-level effects do not materialize by changing the firing rates of neurons but rather develop by modifying synaptic connections.

### Limitations and future directions

A primary limitation of the present study is the absence of direct, *in vivo* measurements of the electric fields generated by the tDCS and tACS currents within the visual cortex. Although quantifying the field strength with depth electrodes would provide the most direct measurement, technical challenges made this unfeasible. However, because our recordings were obtained from the cortical surface directly beneath the stimulation electrodes, the recorded activity is expected to be largely representative of the targeted area and less susceptible to the current shunting or unintended co-activation often associated with protocols targeting deeper structures such as the hippocampus or amygdala.

Furthermore, the parameter space regarding stimulation intensity warrants further exploration. We opted for a protocol strong enough to induce reliable post-stimulation changes while remaining translationally relevant to human safety limits. Prior evidence suggests that weaker fields fail to produce post-stimulation effects (21) and modulate less than one-third of the target neural population (27). Due to the inherent complexities and time constraints of experiments in NHP models, systematically mapping the dose-response curve fell beyond the scope of this investigation; nevertheless, it remains as an interesting question to pursue in future. Specifically, while our findings align with reports of cathodal “excitation” at higher intensities, it remains to be seen whether lower-intensity DC stimulation might yield a “depressive” plasticity effect, as hypothesized in previous literature (15, 77). In a small pilot study, we also studied anodal tDCS but found weaker effects compared to cathodal tDCS and hence pursued with cathodal tDCS. How anodal tDCS affects gamma remains an open question to pursue in future.

Regarding visual stimulation, we utilized full-screen gratings to drive strong, synchronized gamma-band reverberations across the network. However, smaller stimuli are often more effective at driving robust spiking activity within specific receptive fields. Future studies are needed to investigate whether pairing non-invasive neurostimulation with highly preferred, localized visual stimuli alters post-stimulation single-unit firing patterns differently than the wide-field network synchronization observed here.

Finally, due to amplifier saturation and the introduction of hardware noise during stimulation, we were unable to fully eliminate stimulation artifacts from the recordings. Consequently, our analysis is restricted to offline, post-stimulation effects. Future paradigms employing advanced artifact-rejection techniques to resolve online neural activity during the 20-minute stimulation window will be crucial for disentangling the immediate mechanisms of network entrainment from the subsequent consolidation of offline changes.

## MATERIAL & METHODS

### Animal Preparation

The experiments and surgical procedures described here were approved by the Institutional Animal Ethics Committee of the Indian Institute of Science and were in accordance with the guidelines set by the Committee for the Purpose of Control and Supervision of Experiments on Animals (CPCSEA). This study involved two healthy adult bonnet macaques (Macaca radiata): M1, a 15-year-old female weighing 5.8 kg and M2, a 12-year-old male of 7.2 kg body weight. For both monkeys, a titanium headpost was implanted over the frontal region of the skull under general anaesthesia. Four weeks later, after they recovered completely, training for the passive fixation task was started. When the animals learned the task, another surgery was performed to implant a microelectrode array with 6×8 active platinum microelectrodes (48 electrodes, each 1 mm long, with an interelectrode distance of 400 μm, Utah array, Blackrock Neurotech) in the right primary visual cortex. The grid was centred ∼10-15 mm lateral from the midline and ∼10-15 mm rostral from the occipital ridge. Reference wires were either put under the dura or wrapped around the titanium screws near the craniotomy. In both monkeys, another grid of the same size was also implanted in the V4 area, but we did not use that data in this study as the array did not show any visually driven activity in M2. Recording of neural signals from the implant began after two weeks of recovery.

### Experimental Setup

LFP and spike data were recorded when the monkeys sat in a primate chair with their head fixed and performed the passive fixation task. The stimuli were displayed on a monitor (BENQ XL2411, 1280×720 resolution, 100 Hz refresh rate, gamma-corrected) with a mean luminance of 60 cd/m², calibrated using ColorChecker Display Plus. The monitor was placed ∼50 cm away from the eye-level of the subject animals, while eye position data were tracked by the ISCAN ETL-200 Primate tracking system. The monkey and the display setup, along with the eye tracker, were kept in a dark Faraday enclosure with dedicated grounding to minimize interference from external electrical noise. Stimuli were generated and presented pseudo-randomly using custom software.

### Experimental Design and Transcranial Electrical Stimulation

Monkey fixated on a dot at the centre of the screen within a 1.5-degree window around the fixation point. At the beginning of a trial, and after one second of fixation, three stimuli were presented per trial for 800 ms each, with an interval of 700 ms. Full-screen black and white static gratings with a combination of four spatial frequencies (0.5, 1, 2 and 4 cycles/degree), four orientations (0°, 45°, 90°, 135°) and three contrasts (25%, 50% and 100%) were presented to produce strong stimulus-induced gamma activity in primary visual cortex. All the combinations of stimulus features resulted in a block of 48 stimuli, i.e. 16 trials. We ran 9-10 blocks to get a gamma band estimate, which took ∼20 minutes. In total, six gamma protocols were run: Pre-Stimulation, Stimulation, Post-Stimulation, Post+30 min, Post+60 min and Post+90 min. In the stimulation protocol, we applied either current pulses or sham stimulation when the monkey viewed the gratings as in other protocols. To investigate the time course of this effect, post-stimulation gamma protocols were spaced 10 minutes apart.

Transcranial Electrical Stimulation was applied targeting the right primary visual cortex with a two-electrode fronto-occipital montage, resembling the Fz-O2 coordinate of the 10-20 electrode placement system in humans. We used two HD ring electrodes (Soterix Medical) with an outer diameter of 1.2 cm and placed them on the scalp using Ten20 conductive paste (Weaver and Company). For both monkeys, we applied a 3 mA current as cathodal direct current stimulation in our tDCS experiments, which is in line with successful human studies in the past and shown plasticity like changes after stimulation (55) and lower than safe and tolerable intensity of 4 mA as reported in a recent study (78). A previously reported tDCS study on non-human primates employed 2 mA current, but their objective was to study online tDCS effect, and they did not observe any post-stimulation changes (37), so we opted for slightly higher intensity. In tACS experiments, for M1 (Female, head circumference of 22.5 cm), ±1.5 mA current was applied, and in the case of M2 (Male, head circumference of 33cm), ±2.5 mA current was applied. Previously, studies have employed 1.5 mA-2 mA current intensity for shorter duration to monkeys and reported successful changes in spike timing while stimulation was applied (38, 46). Considering the impact of factors such as muscle density and head size on the effect of stimulation (79), the intensity of the current was increased for M2 compared to M1. We applied 20 minutes of current stimulation in each experiment and collected data from multiple sessions (five sessions and three sessions for M1 and M2 in tDCS, and four sessions and three sessions for M1 and M2 in tACS) to replicate our findings.

Sham stimulation experiments were conducted alongside to mimic the sensation of ramping up and down of current at the beginning and end of the protocol, with no stimulation in between. Overall, these recordings were time consuming as the monkeys had to work for more than two hours continuously to finish the entire sequence of protocols. M1 worked for several hours, and hence sham stimulation and actual stimulation protocols could be recorded on the same day (in an alternate order across days). For M2, they were recorded on separate days.

The long experimental duration also prevented us from doing a more systematic analysis of the dose dependence or testing other paradigms such as anodal tDCS. We first ran some pilot tests to find conditions that appeared to sufficiently modulate gamma activity and chose to perform several repetitions of those conditions (typically 4 to 5, along with the same number of sham sessions) to test the robustness of these effects. Several other manipulations that we could not carry out in this work but would be interesting directions to pursue are discussed in the Limitations and Future directions section in the Discussion.

### Data Acquisition and Electrode Selection

Neural data were recorded using the Cerebus neural signal processor (Blackrock Neurotech). The signal obtained was band-pass filtered between 0.3 Hz (analog Butterworth filter, first order) and 500Hz (digital Butterworth filter, fourth order), sampled at 2 kHz and digitized at 16 bits resolution to collect LFP data. Multi-unit activity was recorded by filtering the signals between 250 Hz (digital Butterworth filter, fourth order) and 7.5 kHz (analog Butterworth filter, third order), then applying an amplitude threshold of ±4 (M1) or ±5 (M2) standard deviations of the signal.

Receptive field mapping was done across multiple days by showing a series of small grating stimuli (0.3°-0.4°) of full contrast on an equally spaced locations within a rectangular grid, while the monkey fixated at the centre of the screen. The evoked LFP was fitted with a 2D gaussian to estimate the receptive field centre and spread. The receptive fields with respect to the fixation point were estimated at the eccentricity of 2.6° to 3.3° for M1, and at the eccentricity of 2.8° to 3.5° for M2, located at lower left quadrant of the visual space. Electrodes with stable receptive field centres and impedance values of less than 2500 KΩ were selected for further analysis. According to the cutoffs, we got 32 and 42 microelectrodes from M1 and M2, respectively, which provided LFP data for our experiments. For MUA data, electrodes with a higher signal-to-noise ratio (SNR) and a change in firing rate greater than a preset cutoff (SNR = 1.2 for M1, 1.5 for M2; change in firing rate = 0.3 for M1, 1 for M2) were selected for analysis. The full screen grating stimuli used in the main experiments does not induce strong firing of neurons; thus, the firing rate cutoffs for multi-unit activity were low.

### Data Analysis

Collected data was analyzed using custom codes written in MATLAB (The MathWorks, RRID:SCR_001622). The objective of the study was to understand the post-stimulation effect of transcranial electrical stimulation in terms of stimulus-induced gamma oscillations. Therefore, we focused on pre- and post-stimulation data, estimating power spectral density (PSD), firing rates, and Field-Field coherence. To remove trials with excessive electrical noise or movement-related disruptions, the amplitude of the signal from any trials that exceeded the threshold of six times the standard deviation from the mean signal was deemed noisy and was rejected. For the remaining data, the PSD and time-frequency spectra were computed using the multitaper method with a single taper. The stimulus period was fixed at a range of 250 ms to 750 ms to avoid stimulus onset-related activity. The baseline was kept at -500 ms to 0 ms relative to the stimulus onset. The peak of gamma power increased as the stimulus contrast increased (8). We therefore set the gamma ranges individually for each contrast (Fig. S1).

Gamma band power for each protocol was computed by first averaging across the power values of PSD from a set gamma frequency range, excluding 48-52 Hz (as this range is affected by line noise). We computed the change in gamma power as: Δgamma power = 10*(log (Power_St_) - log (Power_Bl_)), where Power_St_ and Power_Bl_ were calculated by averaging the stimulus and baseline PSD values across the gamma range. Multi-unit firing rate was calculated by binning spikes across the stimulus period of each trial in 10-ms bins and averaging across trials. Coherency was calculated as Coherency_xy_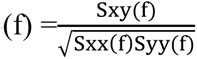. Here, Sxy(f) is the cross-spectrum and Sxx(f) and Syy(f) are the autospectra of each signal. Coherence is the absolute value of coherency. We computed Field-Field coherence using the coherencyc command of the Chronux toolbox, employing the multitaper method with three tapers.

For each monkey, electrodes were pooled across multiple sessions. Statistical test was conducted using t-test between the data obtained from the stimulation session electrodes and the sham session electrodes.

### Model parameters and analysis of simulation output

We utilized a previously reported extension (32) of the Jadi and Sejnowski (JS) model (29) to simulate cortical gamma oscillations in our study. The JS model uses a dual-population (excitatory and inhibitory) network with inhibition-stabilized dynamics to explain how stimulus properties, such as increase in size increases gamma power and increase in contrast increases peak frequency. In the model, population firing rates r_E_ (excitatory) and r_I_ (inhibitory) change according to the following equation, where *f*_E_ and *f*_I_ are sigmoidal activation functions, subscript p denotes population type (E or I) in equation 3.

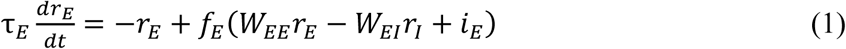

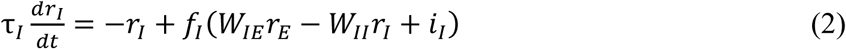

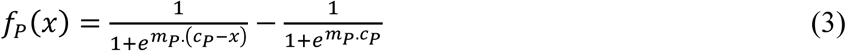

To account for the intermittent, bursty nature of physiological gamma rhythms recorded from the brain, the extended model replaced the steady-state inputs of the original model with Ornstein-Uhlenbeck (OU) noise processes (32).

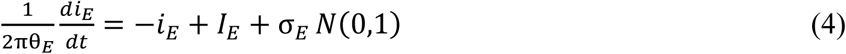

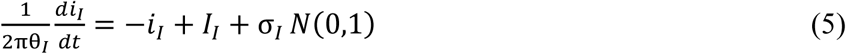

In this formulation, i_E_ and i_I_ represent the time-varying input drives, while I_E_ and I_I_ denote their respective steady-state mean values. Stochasticity is introduced by sampling from a standard Gaussian distribution, *N*(0,1), at each time point. Mathematically, this process is equivalent to passing additive white noise through a first-order low-pass filter with a cutoff frequency θ_E_ or θ_I_ specific to each population.

The following equations from Jadi and Sejnowski’s model ((29); their equation 6), provides analytical approximation of power (which is proportional to σ; see their equation 12), and peak frequency (ω), where *S*_E_ and *S*_I_ are slopes of activation functions.

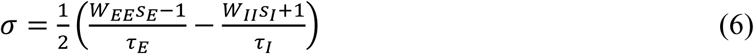

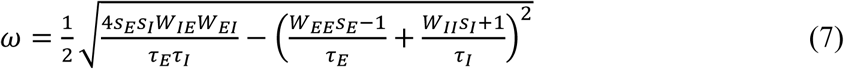

We simulated slow and fast gamma oscillations by adjusting the inhibitory firing rate time constant (*τ_I_*), which determines the central frequency of the generated rhythms. To model slow gamma, we set τ*_I_* to approximately 15.4 ms, representing the kinetics of somatostatin-positive interneurons. For fast gamma, τ*_I_* was reduced to 7.7 ms corresponding to parvalbumin-positive interneurons (See Table S2). These values align with reported time constants for these interneuron types. Additionally, to reflect the lack of recurrent connectivity among SOM cells in V1, the inhibitory-to-inhibitory weight (WII) was set to zero during slow gamma simulations. All other model parameters remained consistent with those detailed in original JS model.

To investigate how synaptic weight adjustments influence gamma power, we simulated proxy LFPs. We began by establishing a super-linear regime, where an increase in inhibitory input (i_I_) drives an increase in oscillation amplitude alongside a decrease in peak frequency. Within this regime, we selected input drive pairs (i_E_ and i_I_) where i_I_ > i_E_ reflecting greater surround activity relative to the receptive field in case of full screen stimuli. We set θ_E_ value at 16, and θ_I_ value at 1, while standard deviation of noise (*_σ_*) was kept same as I_E_ or I_I_, as used in prior models (42).

Each iteration of these simulations was run using forward Euler method (time step: 2×10^−5^ S ) with a time span from -2.047 s to 2.048 s. Mimicking a stimulus display, input drives and variances were set to zero from -2.047 to 0 s, then immediately set to the target values at onset of stimulus presentation, and returned to zero upon completion. While the simulated stimulus duration exceeded that of our empirical experiments, this extended window was chosen to ensure model stability and a steady-state output.

We simulated 15 iterations per combinations and computed proxy LFP as -rE -rI from these iterations and they were passed through fourth order low pass Butterworth filter (cutoff frequency at 200 Hz) and downsampled by a factor of 200, resulting in a sampling frequency of 250 Hz. PSD values were obtained from proxy LFPs by multitaper method using a single taper, as done in actual experimental data. We considered gamma ranges for both slow and fast gamma as ± 10 Hz from respective peak gamma frequency, and this range was considered to estimate average gamma band power for further analysis.

## Acknowledgements

This work was supported by Wellcome Trust/DBT India Alliance (Senior fellowship IA/S/18/2/504003 to S.R.), Institute of Eminence (IoE) funding to IISc, a grant from Pratiksha Trust to IISc (Project REGAIN; IISc-PT-4720-2026) to S.R, and Senior Research Fellowship (SRF) from The University Grant Commission (UGC) awarded to N.M.

## Supplemental Figures

**Fig. S1.**
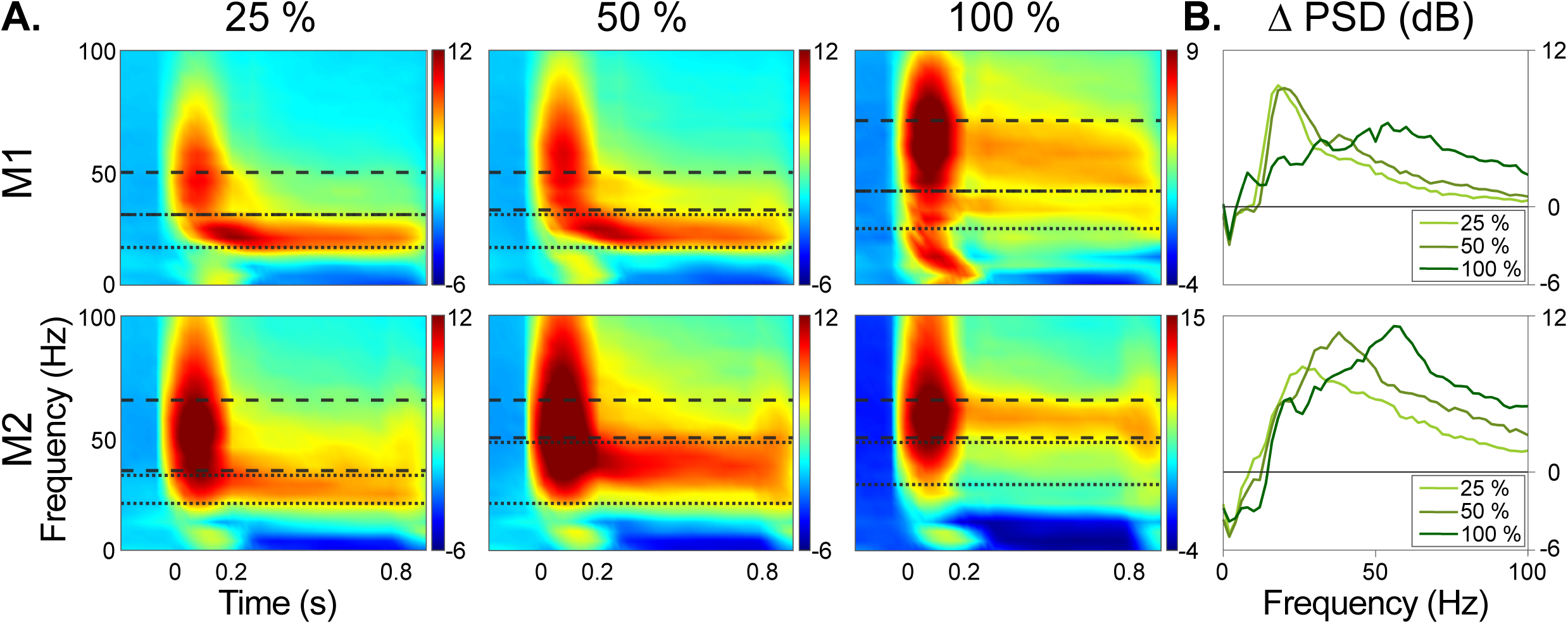
Gamma ranges for stimuli of different contrasts **(A)** Stimulus induced gamma in visual cortex, visualized by time frequency plots for both monkeys. For 25% contrast stimuli, slow gamma ranges are 16 Hz-30 Hz in M1, 20 Hz-32 Hz in M2, fast gamma ranges are 30 Hz-48 Hz in M1, 34 Hz-64 Hz in M2. For stimuli with 50% contrast, slow gamma ranges are 16 Hz-30 Hz in M1, 20 Hz-46 Hz in M2, and fast gamma ranges are 32 Hz-48 Hz for M1, 48 Hz-64 Hz for M2. For 100% contrast, slow gamma ranges are 24 Hz-40 Hz in M1, 20Hz-32 Hz in M2, fast gamma ranges are 40 Hz-70 Hz in M1, and 48 Hz-64 Hz in M2. **(B)** PSD for three different contrast shows contrast dependent distinct gamma profiles, for M1 (up) and M2 (down).

**Fig. S2.**
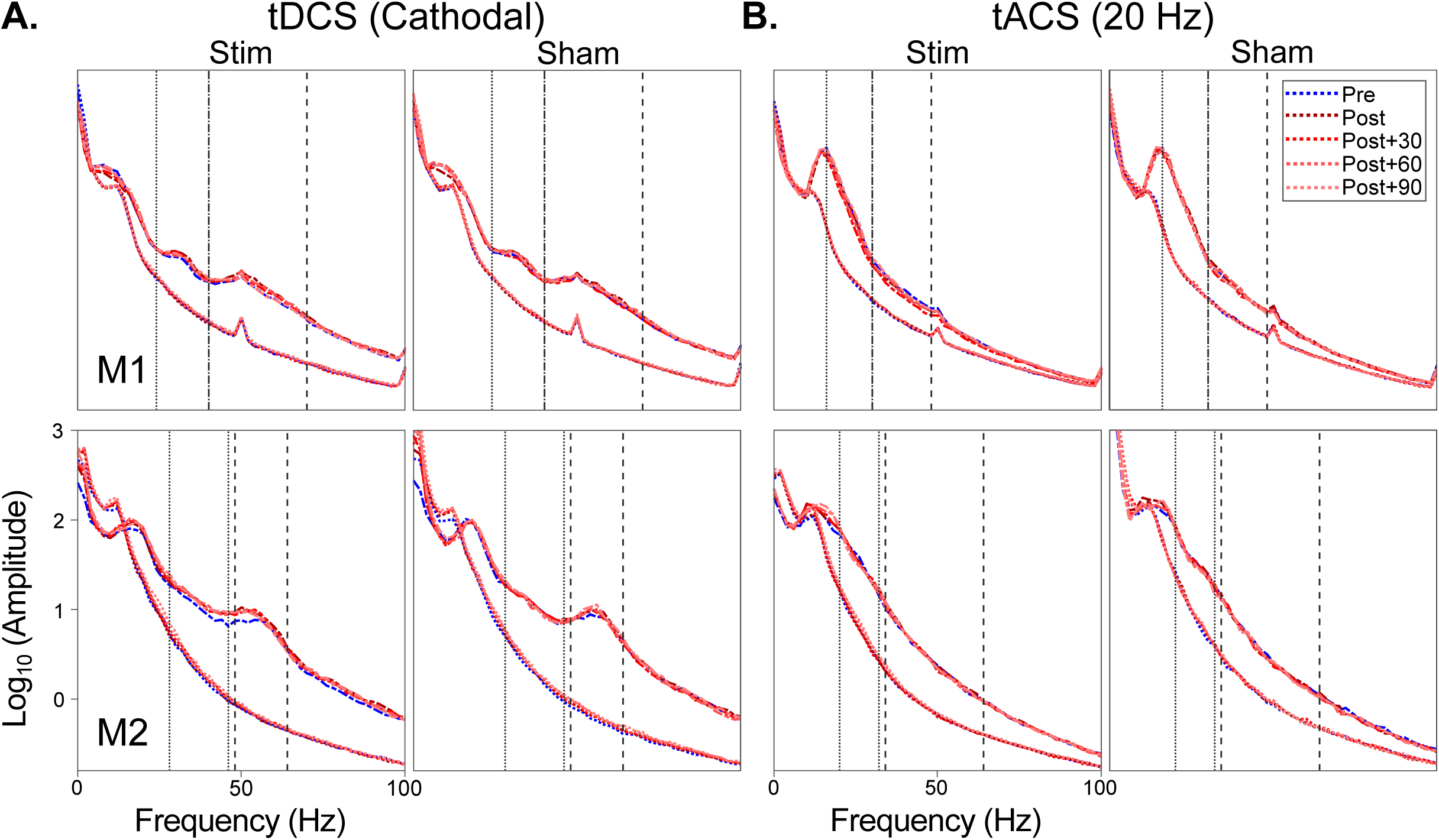
Comparison of baseline powers for both monkeys **(A)** Both baseline (in colored dotted lines), and stimulus period powers (in colored dashed lines) are plotted for tDCS condition with ranges of frequency bands marked. The line for Pre-stim power is plotted in blue. **(B)** Same as **(A)** but for 20 Hz tACS condition.

**Fig. S3.**
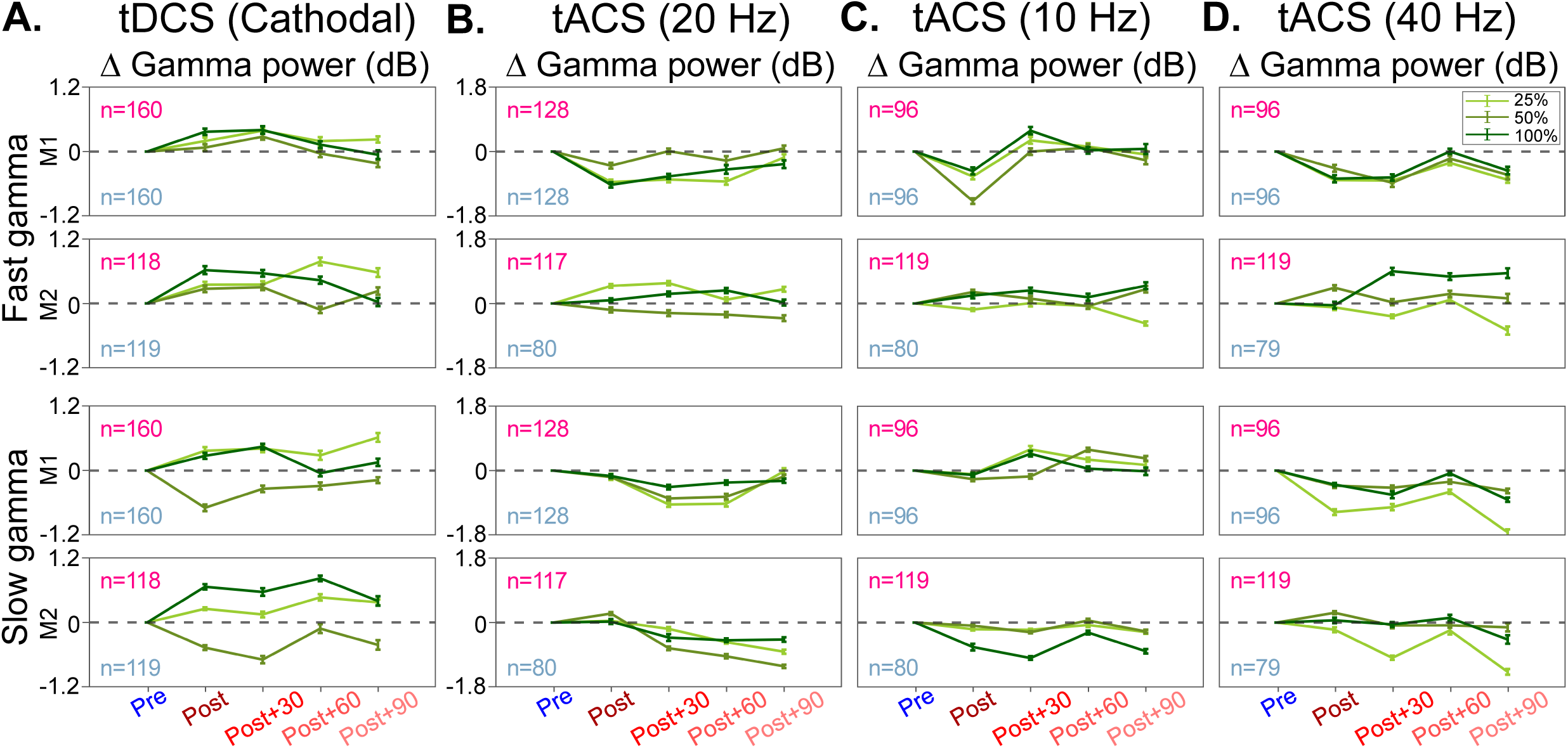
Comparison of contrast specific Effect by tDCS and tACS **(A)** Line plots of different shades of green show comparison of effect of tDCS on stimuli of different contrasts (25%, 50% and 100%) on both the monkeys. Gamma power, averaged across a selected frequency range specific to each contrast, is plotted from all the LFP electrodes pooled from multiple sessions. The higher contrast stimuli show stronger post stimulation effect in increasing gamma band power. **(B)** Effect of 20 Hz tACS in contrast specific manner is plotted in the same format as **(A)**. The lower contrast (25%) stimuli show comparably stronger effects of tACS stimulation. **(C)** Stimulation at 10 Hz shows inconsistent effect across two monkeys. **(D)** tACS at 40 Hz shows suppression of gamma power largely for low contrast (25%) and followed the general trend observed for 20 Hz tACS.

**Fig. S4.**
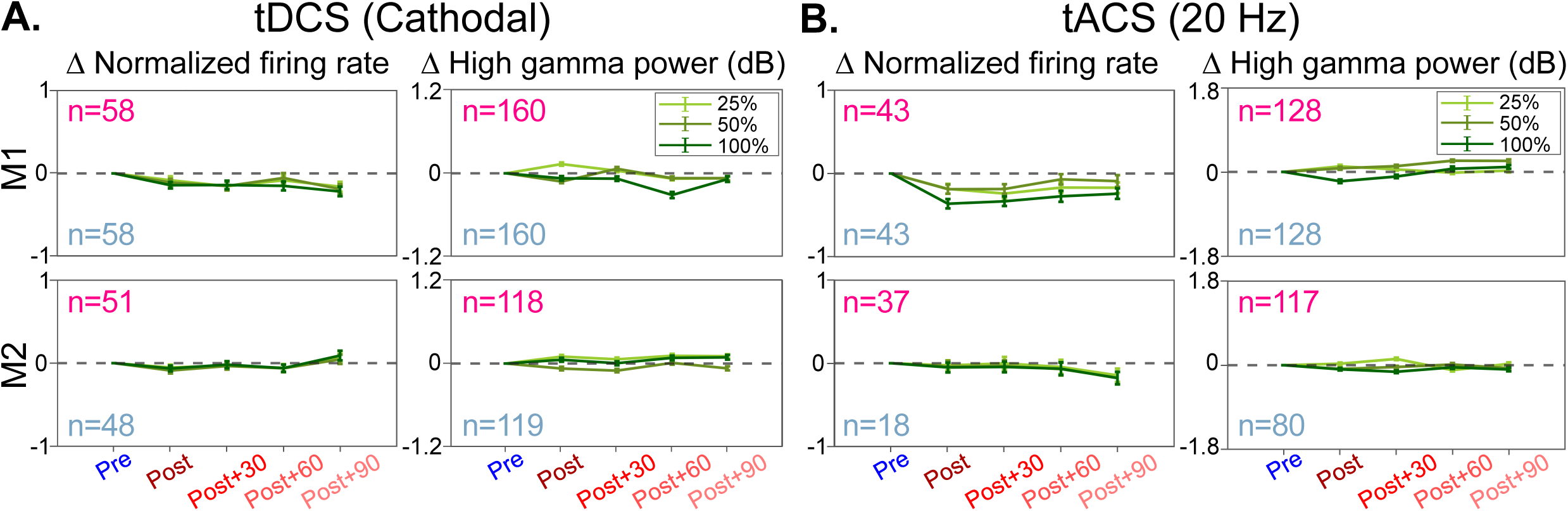
Effect of stimulation on Firing rate and High gamma power **(A, Left panel):** Mean Firing rates are plotted as a line plot with each point corresponding to each protocol, and each shade of green as different contrasts, with data collected from M1 (58 electrodes, stim and sham each), and M2 (51 electrodes from stim and 48 electrodes from sham). **(A, Right panel)**: Average High gamma power (200 Hz-250 Hz) is plotted from all good LFP electrodes across the sessions and different stimulation conditions. **(B)** Same as **(A)**, but for 20 Hz tACS condition.

**Fig. S5.**
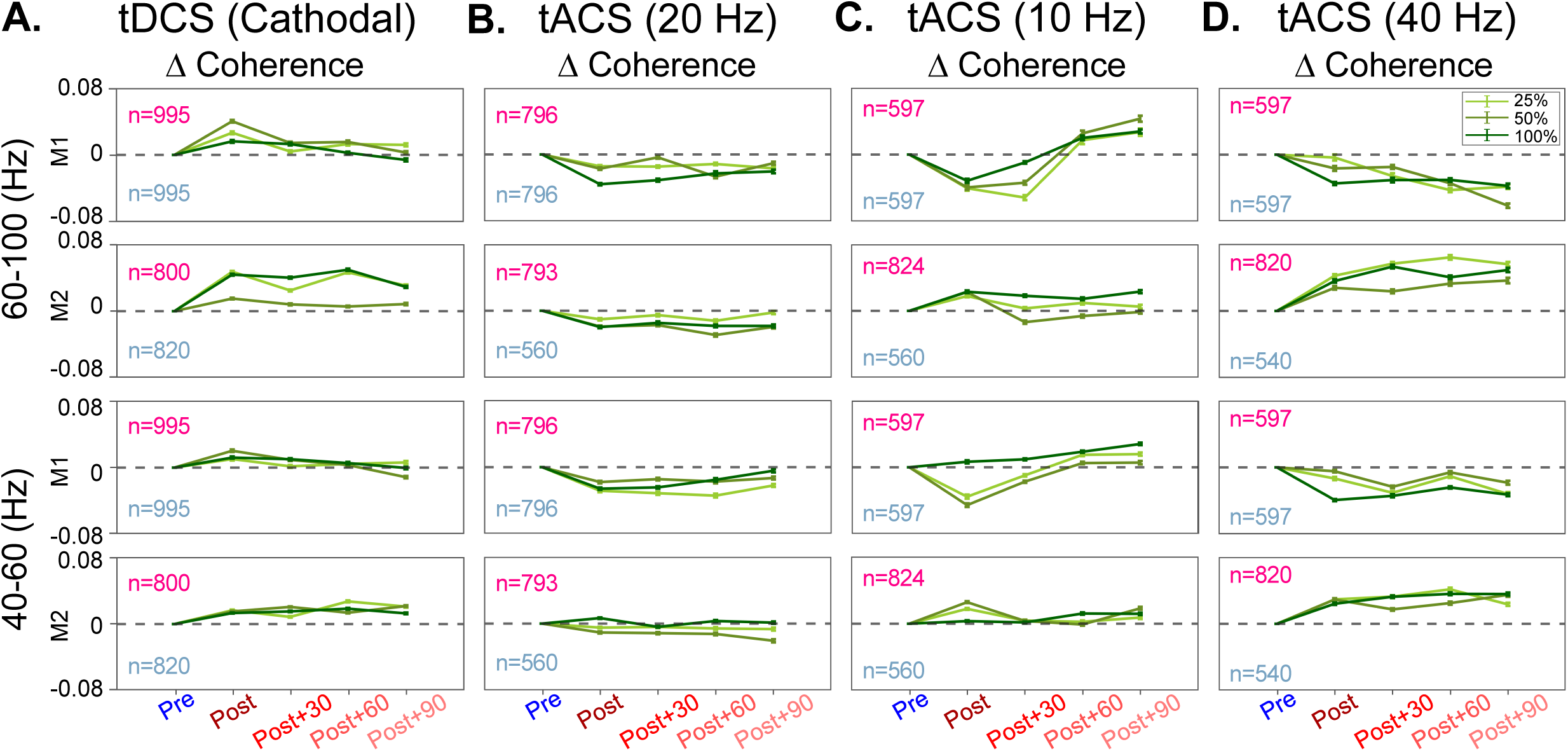
Effect of stimulation on LFP-LFP Coherence for different contrasts **(A)** Effect of cathodal tDCS plotted for stimuli of 25%, 50% and 100% contrast by line plots of different shades of green. Coherence values are calculated for a specific frequency range (60 Hz-96 Hz for M1, and 66 Hz-96 Hz for M2) and averaged across electrode pairs within the 0.4 mm-1.2 mm range. Pre-stimulation coherence values were subtracted from all post-stimulation coherence values to generate the line plot. For lower panels, the same was done for 40 Hz-60 Hz range. **(B)** Effect of tACS at 20 Hz on LFP-LFP coherence plotted as line plots are compared for different contrasts. The calculation of mean coherence is done on the frequency range of 40 Hz-60 Hz for all the conditions. **(C)** Same as **(B)** but for 10 Hz tACS. **(D)** Same as **(B)** but for 40 Hz tACS.

**Fig. S6.**
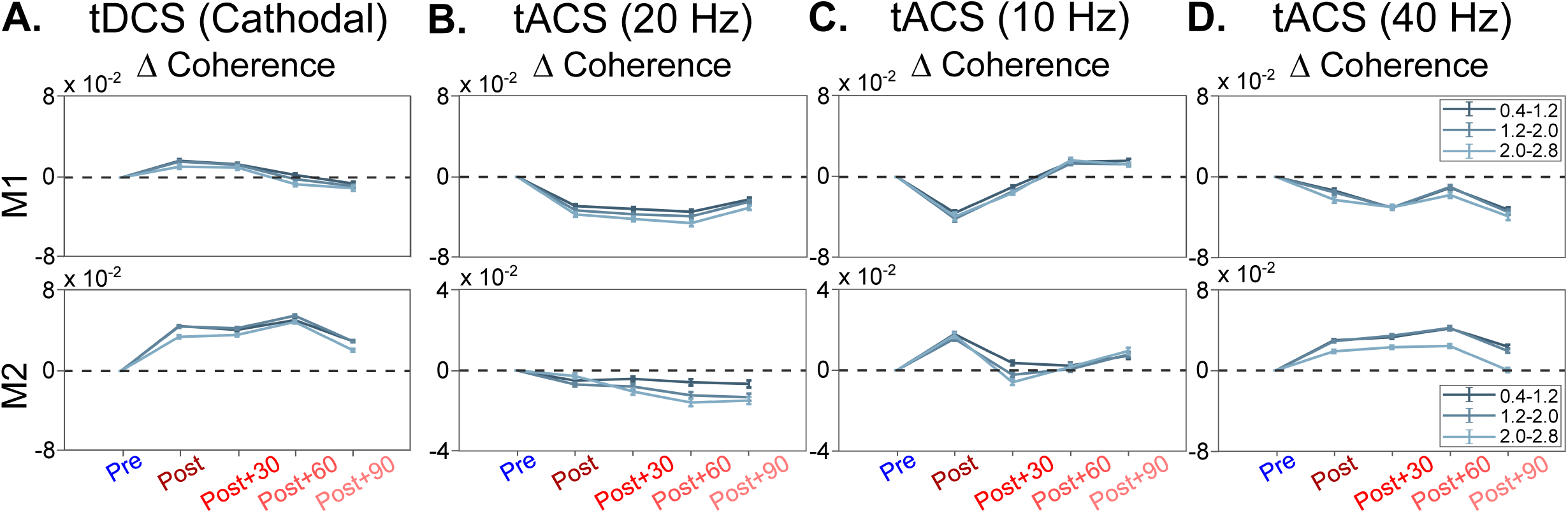
Impact of inter-electrode distance on stimulation-induced change in LFP-LFP coherence **(A)** Different shades of grey denoted three ranges while computing LFP-LFP coherence, 0.4 mm-1.2 mm, 1.2 mm-2.0 mm and 2.0mm-2.8 mm of inter-electrode distance. For cathodal stimulation condition, we have plotted results from 100% contrast and delta coherence values were computed by subtracting pre-stimulation coherence values from all post stimulation coherence values. **(B)** For tACS at 20 Hz, data is shown for 25% contrast, calculated in the same manner as **(A)**. **(C)** Same as **(B)** but for 10 Hz tACS. **(D)** Same as **(B)** but for 40 Hz tACS.

**Fig S7.**
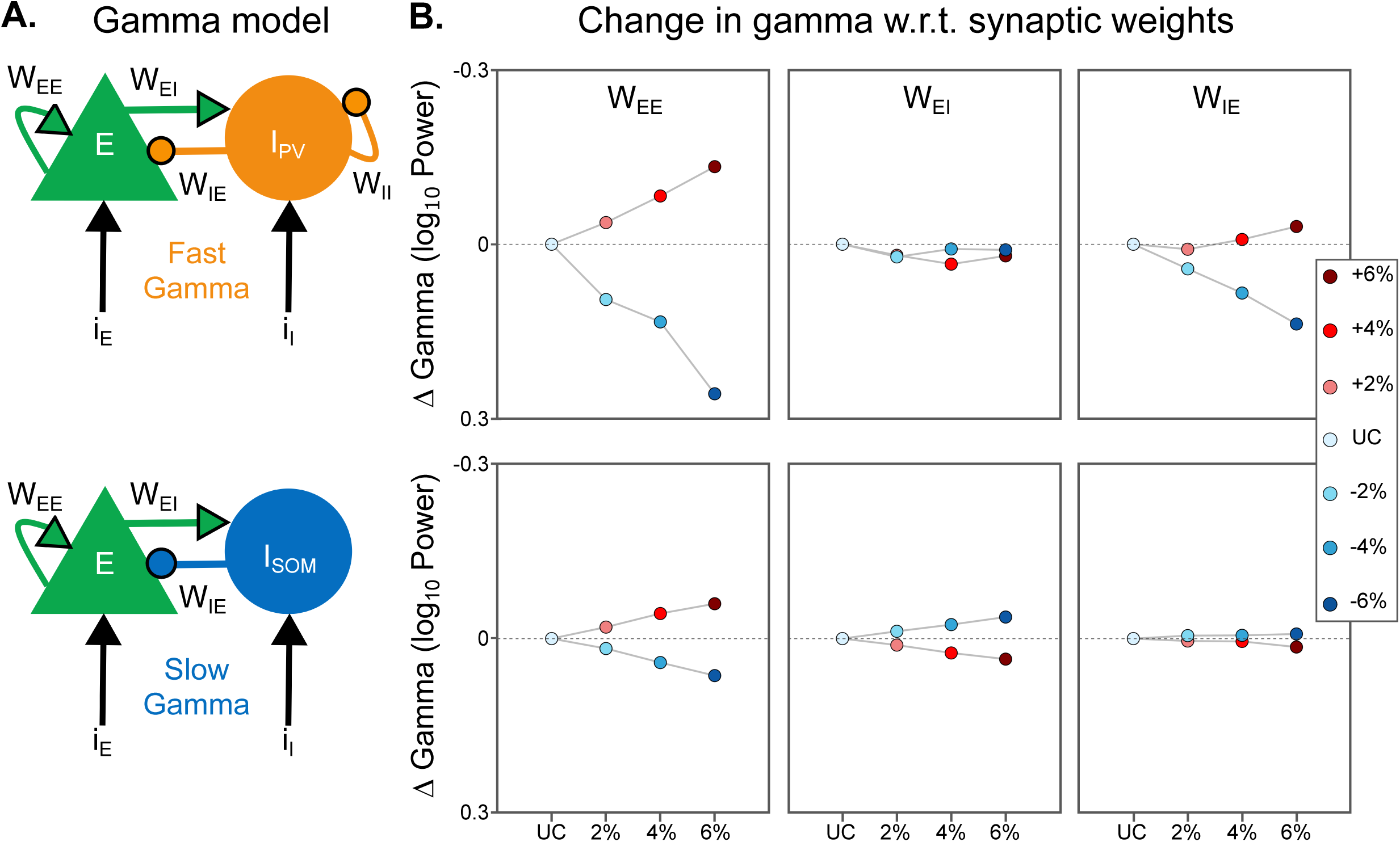
Gamma model and change in gamma power with changes in synaptic weights **(A)** Model of gamma oscillations. Fast gamma model (top) with Pyramidal neuron (E) and Parvalbumin positive PV+ interneuron (I_PV_) and Slow gamma model (bottom) with Somatostatin positive SST+ interneuron (I_SOM_). **(B)** The Averaged gamma power subtracted from the unchanged condition is plotted with percent changes in synaptic weight in the respective columns. Increase in weights is shown in increasing saturation of reds, while decreasing weights is shown in increasing saturation of blues. The top row corresponds to fast gamma, and bottom row corresponds to Slow gamma simulations.

**Fig. S8.**
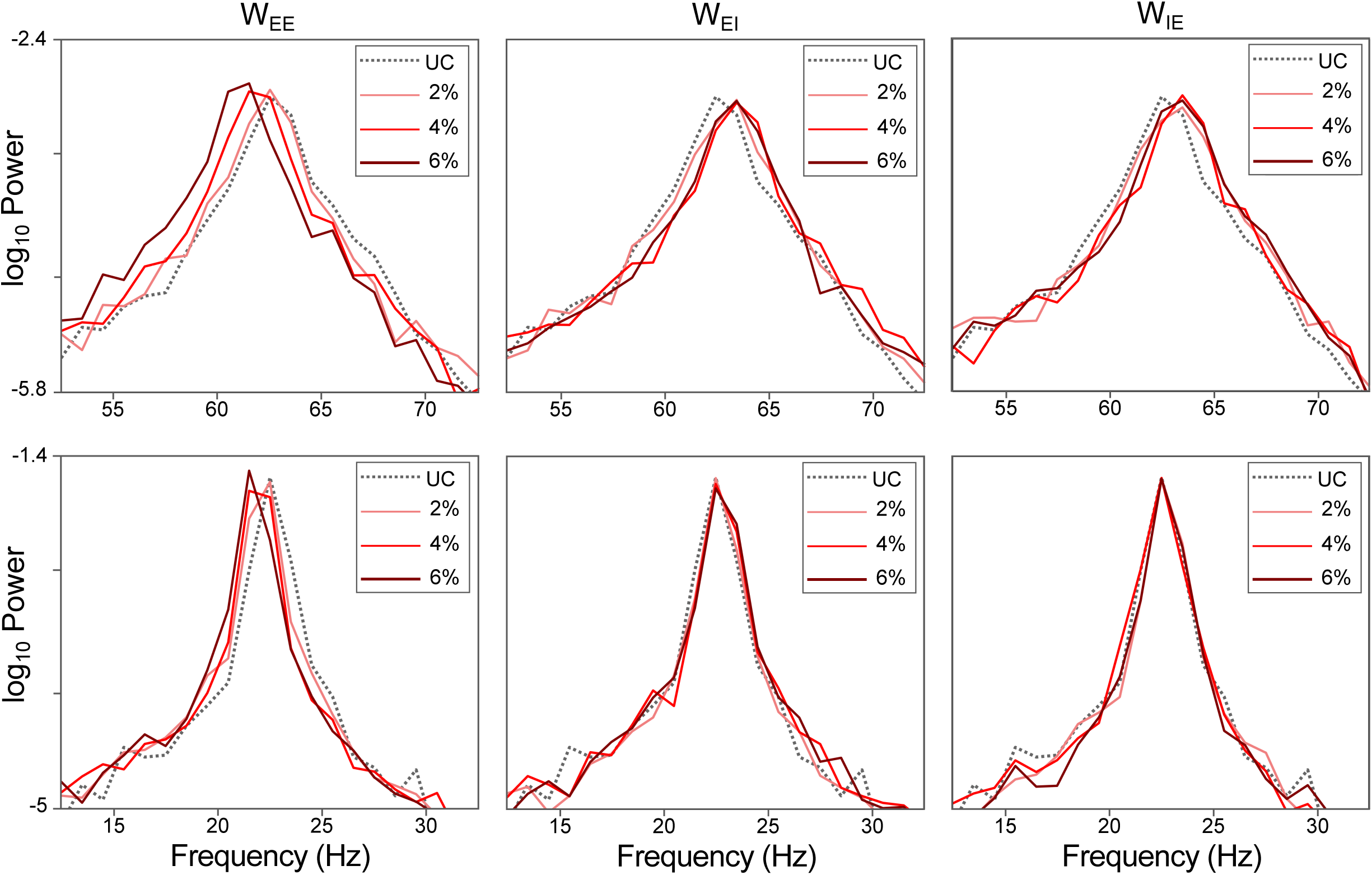
Simulated PSD for increase in synaptic weights in gamma model **(A)** W_EE_, W_EI_ and W_IE_ were increased separately in fast gamma model, and simulated PSD shows trends of change in gamma power and gamma centre frequency, resembling analytical results reported previously. **(B)** Same as **(A)** but for slow gamma model.

**Fig. S9.**
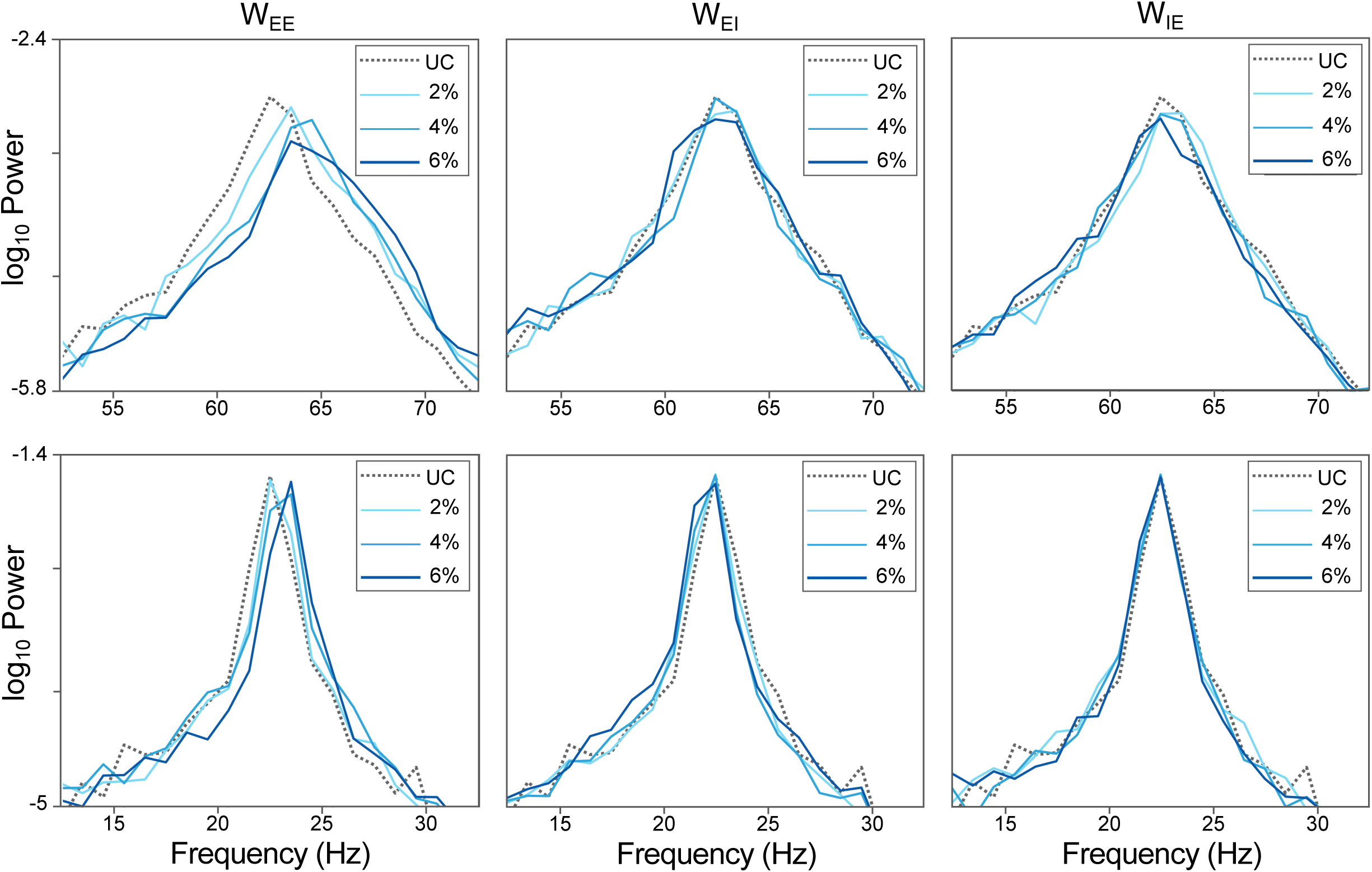
Simulated PSD for decrease in synaptic weights in gamma model **(A)** Synaptic weights were decreased individually, and simulated PSDs show change in gamma power and frequency. **(B)** Same as **(A)** but for slow gamma model.

**Fig. S10.**
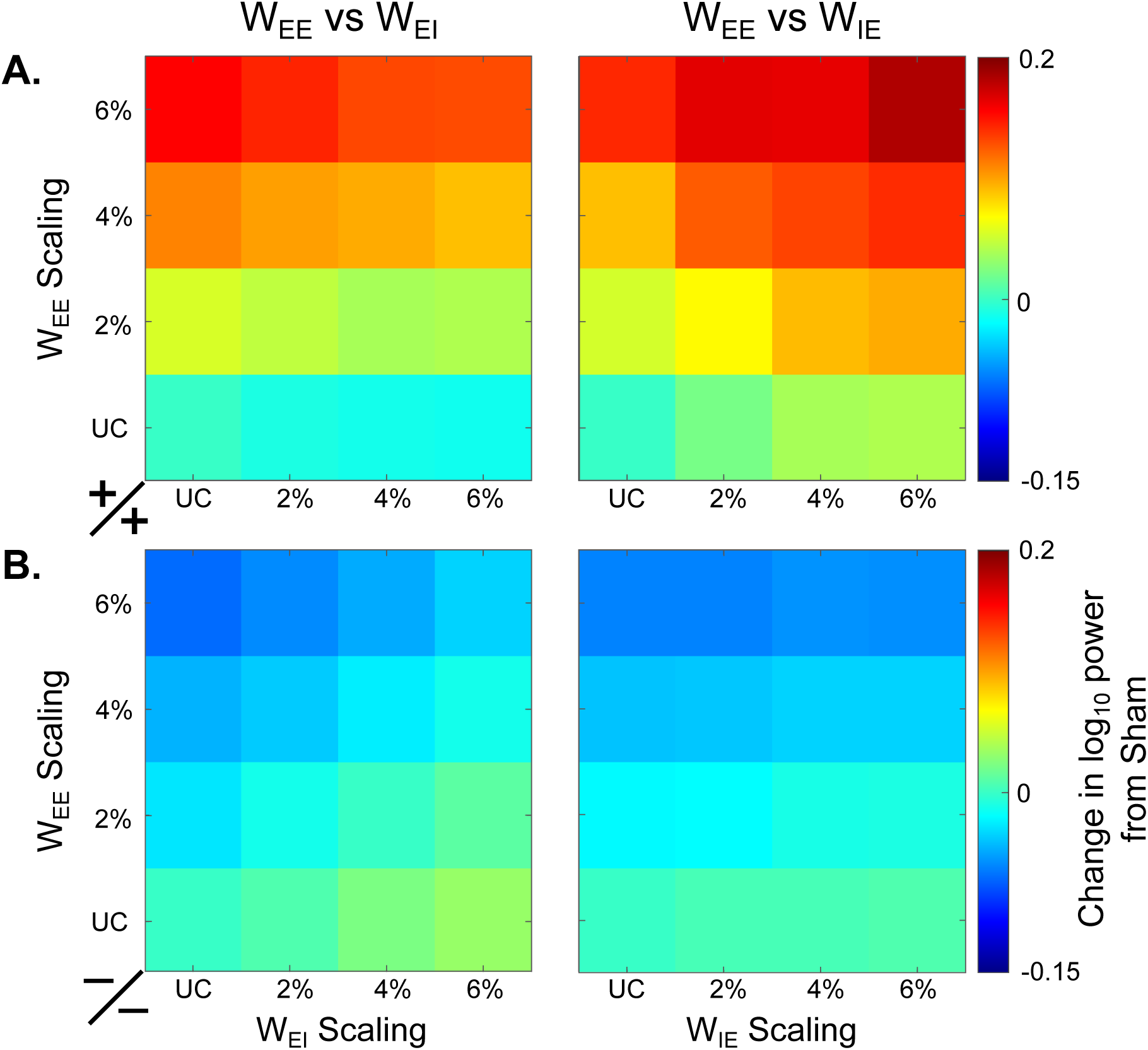
Evolution of gamma power while both W_EE_ and W_EI_ (first column) and W_EE_ and W_IE_ (second column) were changed incrementally. **(A)** Mirroring tDCS condition on fast gamma model with increase in synaptic weights, averaged gamma power plotted in each tile was subtracted from the sham or unchanged condition. Color code in heatmap shows change in gamma power. **(B)** For tACS condition on slow gamma model, and incremental decrease in synaptic weights, calculated in the same manner as **(A)**.

---

**Table S1.** Pre stimulation Gamma power was first subtracted from all post stimulation blocks to eliminate any session specific variability, then between two conditions- stim and sham, paired two sample t test was conducted. For both tDCS and tACS, and both monkeys (M1 and M2) p values and corresponding t values are shown. Number of LFP electrodes across the session is mentioned within bracket along with subject name. P values of significantly different protocols are marked with asterix, and number of asterix shows the significance level, as p<0.05*, p<0.005**, p<0.0005***.

| Stimulation type | Subject Name |  | Protocols |  |  |  |
| --- | --- | --- | --- | --- | --- | --- |
|  | Fast Gamma |  | Post | Post+30 | Post+60 | Post+90 |
| **tDCS**<br>(Fast Gamma) | M1 (n=160) | p = | 1.26x10<sup>-9</sup> *** | 1.71x10<sup>-8</sup> *** | 0.043 * | 0.467 |
|  |  | t = | 6.26 | 5.78 | 2.03 | -0.73 |
|  | M2 (n=118) | p = | 1.40x10<sup>-14</sup> *** | 2.61x10<sup>-17</sup> *** | 5.9x10<sup>-10</sup> *** | 0.738 |
|  |  | t = | 8.35 | 9.21 | 6.46 | 0.33 |
| **tDCS**<br>(Slow Gamma) | M1 (n=160) | p = | 3.13x10<sup>-6</sup> *** | 1.08x10<sup>-15</sup> *** | 0.39 | 0.026 * |
|  |  | t = | 4.76 | 8.47 | -0.84 | 2.23 |
|  | M2 (n=118) | p = | 1.29x10<sup>-29</sup> *** | 1.579x10<sup>-13</sup> *** | 3.45x10<sup>-36</sup> *** | 9.016x10<sup>-6</sup> *** |
|  |  | t = | 13.239 | 8.08 | 15.61 | 4.56 |
| **tACS**<br>(Fast Gamma) | M1 (n=128) | p = | 8.32x10<sup>-27</sup> *** | 1.11x10<sup>-20</sup> *** | 4.57x10<sup>-18</sup> *** | 0.066 |
|  |  | t = | -12.40 | -10.19 | -9.379 | -1.846 |
|  | M2 (n=117) | p = | 9.80x10<sup>-17</sup> *** | 1.32x10<sup>-17</sup> *** | 0.126 | 1.86x10<sup>-8</sup> *** |
|  |  | t = | 9.13 | 9.62 | 1.536 | 5.876 |
| **tACS**<br>(Slow Gamma) | M1 (n=128) | p = | 0.037 * | 9.35x10<sup>-34</sup> *** | 1.11x10<sup>-23</sup> *** | 0.806 |
|  |  | t = | -2.09 | -14.11 | -11.5 | -0.24 |
|  | M2 (n=117) | p = | 0.31 | 0.0007 ** | 1.38x10<sup>-23</sup> *** | 1.98x10<sup>-30</sup> *** |
|  |  | t = | 1.00 | -3.45 | -11.93 | -14.54 |

**Table S2.**
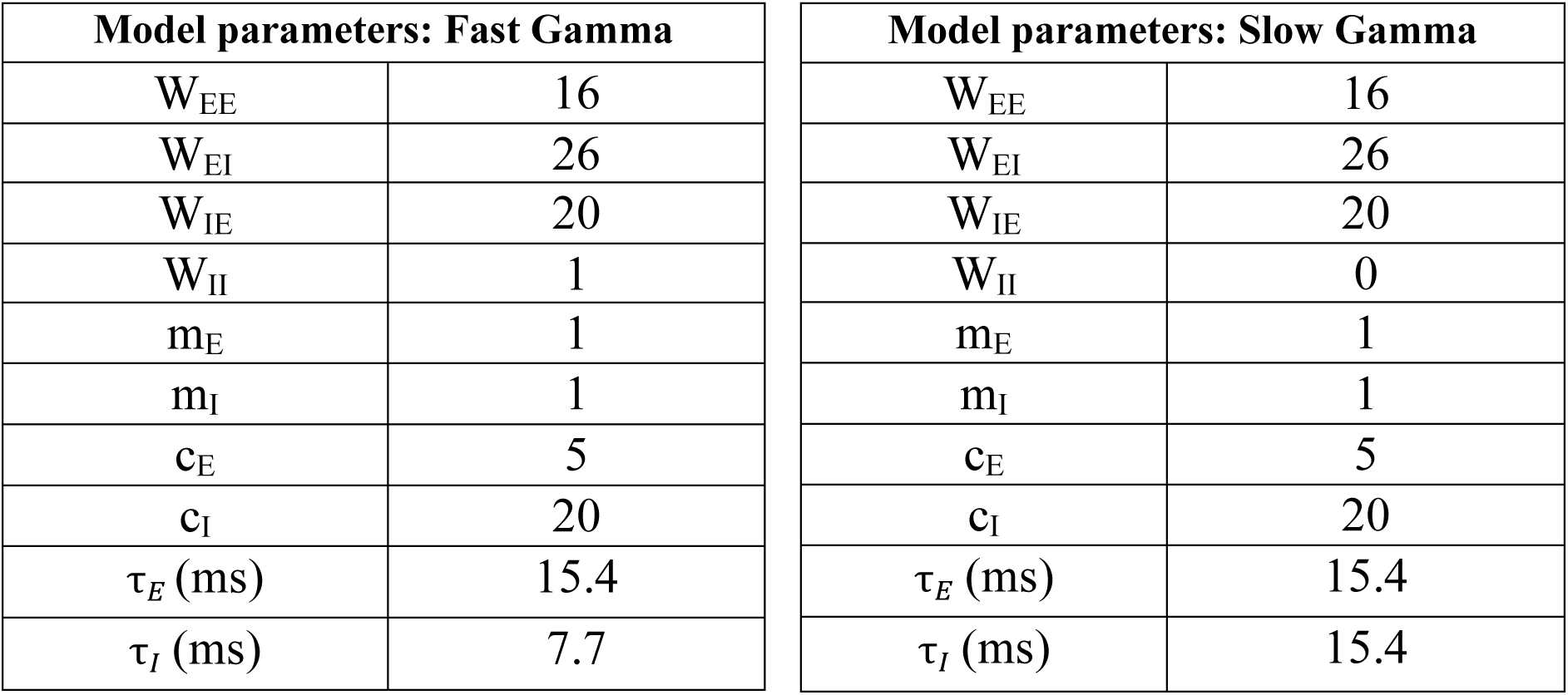
Model parameters used in simulation of gamma oscillation. Fast gamma (left) and slow gamma (right) was modelled by changing the τ*_I_* value. All these parameter values have been used in previous studies modelling stimulus induced gamma oscillation (32, 42).

